# Dietary propionate induces intestinal oxidative stress via inhibition of SIRT3-mediated SOD2 depropionylation

**DOI:** 10.1101/2020.08.10.245399

**Authors:** Qian-wen Ding, Zhen Zhang, Yu Li, Hong-liang Liu, Qiang Hao, Ya-lin Yang, Einar Ringø, Rolf Erik Olsen, Jihong Liu Clarke, Chao Ran, Zhi-gang Zhou

## Abstract

Propionate is a commonly used preservative in various food and feedstuffs and has been regarded as a food additive without safety concerns. However, we observed that dietary propionate supplementation induced intestinal damage in the context of high fat diet (HFD) in zebrafish. The intestinal damage was attributable to oxidative stress owing to impaired antioxidant capacity, which was caused by compromised SOD2 activity in the intestine. Global lysine propionylation analysis of the intestinal samples showed that SOD2 was propionylated at K132, and further biochemical assays demonstrated that K132 propionylation suppressed SOD2 activity. In addition, SIRT3 could directly interact with SOD2 and played an important role in regulating SOD2 activity via modulating depropionylation, and the enhanced SOD2 propionylation in zebrafish fed high fat plus propionate diet was attributable to reduced SIRT3 expression. Finally, we reveal that intestinal oxidative stress resulting from SOD2 propionylation contributed to the compositional change of gut microbiota, which further deteriorated intestinal oxidative stress independent of SIRT3. Collectively, the results in this study reveal a link between protein propionylation and intestine health, and suggest potential risk of a widely used food preservative in HFD context.

## Introduction

Propionic acid (PPA) is a ubiquitous short-chain fatty acid (SCFA), and is a major fermentation product of the enteric microbiome (Koh et al., 2016). As an anti-bacterial compound, propionate is one of the most commonly used preservatives with a maximum allowed concentration up to 0.5% in various foods, such as in cheeses and baked goods, and in animal feedstuffs (Rose, 2013). Propionate inhibits bacterial growth via interrupting enzyme activity and DNA replication (Ng and Koh, 2017). Although propionate is regarded as a food additive without safety concerns (Rose, 2013), several studies have indicated that exposure to propionate may cause mitochondrial dysfunction (Matsuishi et al., 1991; Pougovkina, 2016; Stumpf et al., 1980). Furthermore, studies of autism spectrum disorders (ASD) showed that overproduction of propionate resulting from enriched propionate-producing bacteria in individuals with ASD are potentially toxic to the mitochondria (Frye et al., 2015).

As a SCFA, propionate crosses the mitochondrial inner membrane and serves as precursor for generation of propionyl-CoA, which could enter the tricarboxylic acid (TCA) cycle for energy metabolism or act as a propionyl-CoA donor for lysine propionylation (Schonfeld & Wojtczak, 2016; Flavin & Ochoa, 1957; Chen et al., 2007; Cheng et al., 2009). Lysine propionylation is a common post-translational modification (PTM) existing in histones of eukaryotic cells, such as 293T cells and yeast (Chen et al., 2007; Cheng et al., 2009; Liu et al., 2009; Zhang et al., 2009). Similar to acetylation and butyrylation, histone propionylation is a marker of active chromatin (Kebede et al., 2017). In contrast to histone propionylation, reports about non-histone propionylation in eukaryotic cells are scarce. Cheng et al. reported the presence of lysine propionylation in three non-histone proteins in 293T cells, i.e., p53, p300, and CREB-binding protein (Cheng et al., 2009). Propionate exhibits mitochondrial toxicity and inhibits mitochondrial respiration in liver and muscle due to significant propionyl-CoA accumulation (Matsuishi et al., 1991). Impaired mitochondrial respiration leads to enhanced production of ROS (Bhatti et al., 2017). Imbalance between ROS generation and clearance accounts for oxidative stress, which leads to mitochondrial dysfunction (Bhatti et al., 2017; Wei et al., 1998; Duchen, 2004; Pieczenik & Neustadt, 2007). Recently, a study phenocopying propionyl-CoA carboxylase deficiency suggested a direct connection between propionyl-CoA accumulation and mitochondrial dysfunction caused by protein propionylation (Pougovkina, 2016). However further identification of propionylated proteins resulting in oxidative stress is insufficient.

Studies have demonstrated that gastrointestinal (GI) diseases are associated with ROS and mitochondrial dysfunction. Oxidative stress has important pathogenetic implications for inflammatory bowel disease (IBD) (Palucka, 2007), and enterocytes with abnormal mitochondrial structure have been reported in IBD patients (Novak & Mollen, 2015). The GI tract injury effect of nonsteroidal anti-inflammatory drugs (NSAID) is associated with disruption of mitochondrial structure and function (Rafi, 1998; Somasundaram, 1997; Kyle, 2014). Similarly, dextransodiumsulfate (DSS) induces ROS-mediated inflammation in human colonic epithelial cells (Bhattacharyya et al., 2009). On the other hand, antioxidant drugs, such as sulfasalazine, have shown beneficial effects in the treatment of IBD (Bhattacharyya et al., 2014).

The intestinal epithelium is prone to oxidative damage induced by luminal oxidants because it locates at the interface between an organism and its luminal environment (Circu & Aw, 2012). In this study, we observed that propionate induced oxidative damage to zebrafish intestine in the context of high fat diet. We revealed a mechanism for propionate-induced intestinal oxidative damage that involved propionylation. Superoxide dismutase 2 (SOD2) can be propionylated at the lysine 132 site, which suppressed its activity and resulted in oxidative damage in the intestine. Furthermore, we found that the higher propionylation of SOD2 was due to reduced intestinal expression of SIRT3 in zebrafish fed high fat plus propionate diet. In addition, the intestinal microbiota induced by high fat plus propionate diet also contributed to intestinal oxidative stress, in a SIRT3-independent manner.

## Results

### Propionate supplementation in high fat diet induces intestinal damage

We established a propionate-feeding model via feeding one-month old zebrafish either low-fat diet (LFD), low-fat diet supplemented with 0.5% sodium propionate (LFSP0.5), high-fat diet (HFD) or high-fat diet supplemented with 0.5% sodium propionate (HFSP0.5) (Supplementary Table 1). Although both oil red staining of liver sections and hepatic TG quantification in zebrafish fed HFSP0.5 diet showed lower lipid accumulation (Supplementary Fig. 1A and 1B), histopathologic analysis of H&E-stained intestine sections showed damage (i.e., breaches in the intestinal epithelium and injury to or loss of intestinal villi) in zebrafish fed HFSP0.5 diet (Fig. 1A). The activation of intestinal caspase-9 (Fig. 1B), caspase-6 (Fig. 1C) and caspase-3 (Fig. 1D) was observed in zebrafish fed HFSP0.5 diet, suggesting that a mitochondrial pathway of apoptosis was activated by the HFSP0.5 diet. Meanwhile, intestinal caspase-8 and caspase-12 activity in zebrafish fed HFSP0.5 diet was similar to that in zebrafish fed HFD (Supplementary Fig. 1C and 1D). However, it is worth noting that sodium propionate supplementation in the context of LFD did not induce damage (Fig. 1A) or the elevation of intestinal caspase-9 and caspase-3 activity (Fig. 1E and 1F), which indicated that sodium propionate becomes a damage-inducing factor in the context of HFD.

**Fig. 1.**
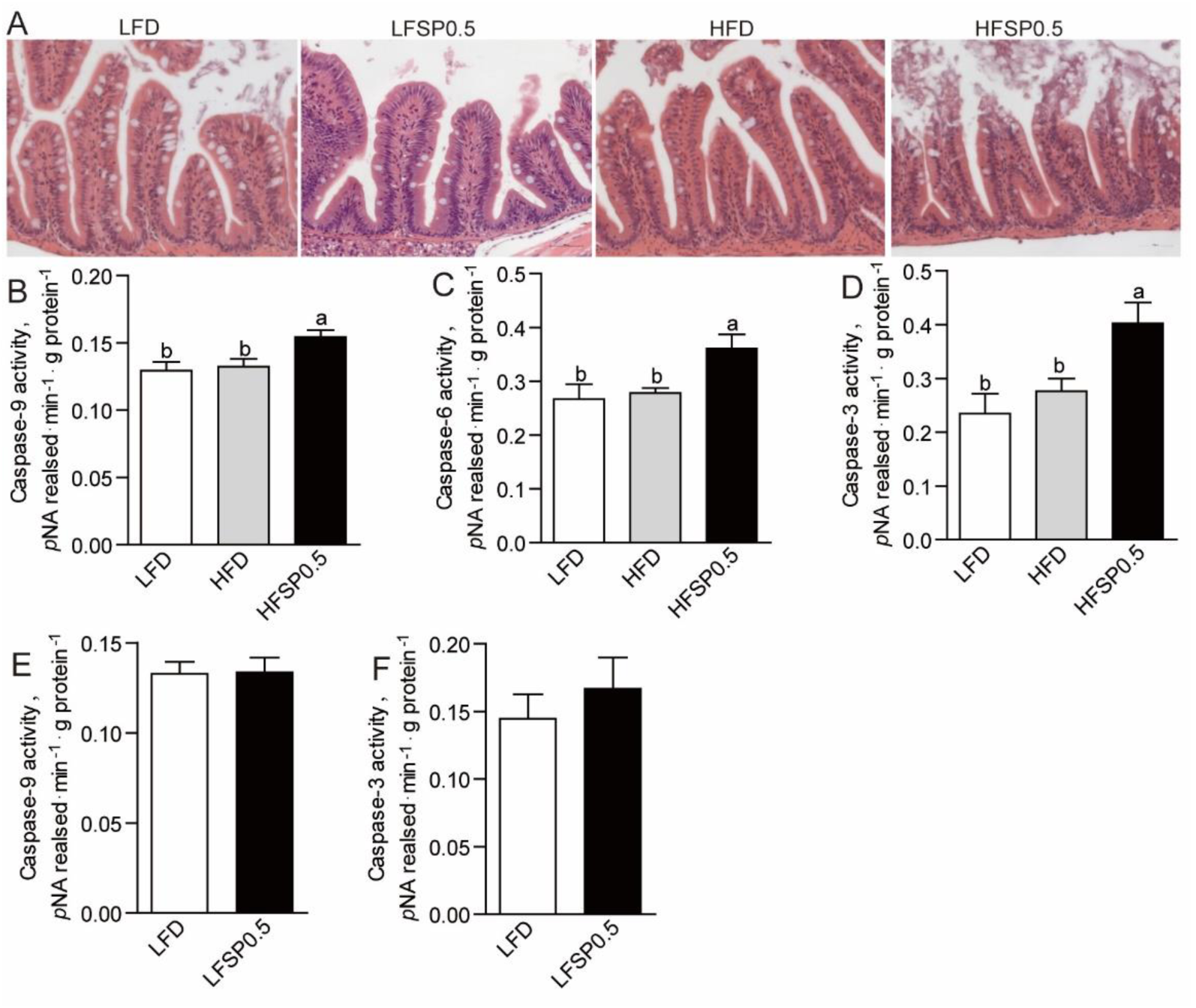
Propionate induces intestinal damage in the context of high fat diet. (A) Representative histopathologic image of H&E-stained intestine sections. (B) Caspase-9, (C) caspase-6 and (D) caspase-3 activity in the intestine of 1-month-old zebrafish fed LFD, HFD or HFSP0.5 diet for 2 wks. (E) Caspase-9 and (F) caspase-3 activity in the intestine of 1-month-old zebrafish fed LFD or LFSP0.5 diet. Values are means ± SEMs (*n*=4∼6 biological replicates). Means without a common letter are significantly different, *P*<0.05. LFD, low-fat diet; HFD, high-fat diet; HFSP0.5, high-fat diet supplemented with 0.5% sodium propionate; LFSP0.5, low-fat diet supplemented with 0.5% sodium propionate.

### Propionate induces intestinal oxidative stress in zebrafish fed high fat diet

To characterize intestinal damage caused by HFSP0.5 diet, we analyzed the difference in mitochondrial membrane potential (MMP) between zebrafish fed LFD, HFD and HFSP0.5 diet. Compared to zebrafish fed HFD, zebrafish fed HFSP0.5 diet showed significant decrease in intestinal MMP (Fig. 2A), which indicated that HFSP0.5 diet caused mitochondrial dysfunction. HFSP0.5 diet caused oxidative stress in zebrafish intestine, as shown by levels of mitochondrial reactive oxygen species (ROS) (Fig. 1B), malonaldehyde (MDA) (Fig. 2C), and protein carbonyl (PC) content (Fig. 2D), whereas LFSP0.5 diet exerted no effect on mitochondrial ROS (Fig. 2E). Furthermore 4-Hydroxy-TEMPO, a membrane-permeable radical scavenger, alleviated the damage induced by HFSP0.5 diet to intestinal epithelium (Fig. 2F and 2G). We treated the ZF4 cell line with a mixture of 150 μM oleic acid and 50 μM palmitic acid (OPA), or a mixture of 150 μM oleic acid, 50 μM palmitic acid and 50 mM sodium propionate (OPP) to mimic the intestinal damage found with the HFSP0.5 diet. The OPP treatment resulted in 18.5% decrease in cell survival rate (Fig. 2H) and 25.2% increase in cell apoptotic rate (Fig. 2I) at 24 h. OPP exposure resulted in 10.8% increase in cellular ROS at 24 h (Fig. 2J).

**Fig. 2.**
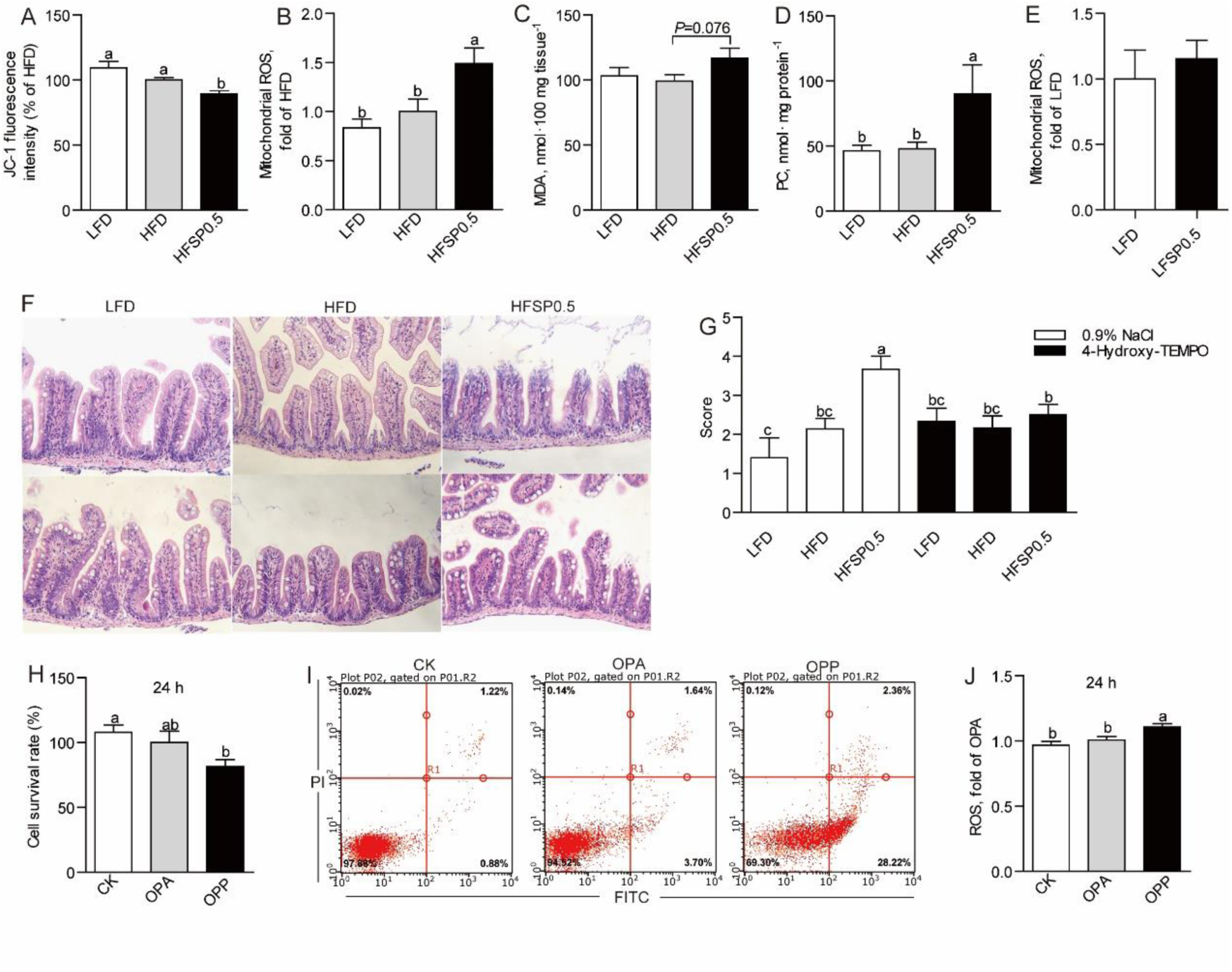
Propionate induces intestinal oxidative stress in the context of high fat diet. (A) The mitochondrial membrane potential in the intestine of 1-month-old zebrafish fed LFD, HFD or HFSP0.5 diet for 2 wks. (B-D) Intestinal biomarkers for oxidative stress in 1-month-old zebrafish fed LFD, HFD or HFSP0.5 diet for 2 wks, including (B) mitochondrial ROS, (C) MDA and (D) PC. (E) Mitochondrial ROS in the intestine of 1-month-old zebrafish fed LFSP0.5 diet. (F) Representative histopathologic images of H&E-stained intestine sections in zebrafish intraperitoneally injected with 4-Hydroxy-TEMPO, a membrane-permeable radical scavenger. (G) Histological score measuring the severity of the intestinal damage of zebrafish intraperitoneally injected with 4-Hydroxy-TEMPO. (H) Cell survival rate and (I) cell apoptotic rate in ZF4 cells treated with a mixture of OPA or OPP for 24 hrs. (J) Cellular ROS in ZF4 cells treated with a mixture of OPA or OPP for 24 hrs. Values are means ± SEMs, for A-D *n*=5 or 6 biological replicates, for E *n*=3 or 4 biological replicates, for G and H *n*=5∼12 biological replicates, for J *n*=8 biological replicates. Means without a common letter are significantly different, *P*<0.05. ROS, reactive oxygen species; MDA, malonaldehyde; PC, protein carbonyl; OPA, mixture of 150 μM oleic acid and 50 μM palmitic acid; OPP, mixture of 150 μM oleic acid, 50 μM palmitic acid and 50 mM sodium propionate.

### Propionate inhibits intestinal total antioxidant capacity in the context of high fat diet

Compared with zebrafish fed HFD, zebrafish fed HFSP0.5 diet displayed lower total antioxidant capability (T-AOC) (Fig. 3A) and SOD2 activity (Fig. 3B). However, there was no difference in the activity of other intestinal antioxidant enzymes, such as glutathione peroxidase (GPx) (Fig. 3C) and catalase (CAT) (Fig. 3D) between zebrafish fed HFD and HFSP0.5 diet. These results suggested that inhibition of SOD2 activity induced by HFSP0.5 diet mainly contribute to impaired T-AOC and oxidative stress.

**Fig. 3.**
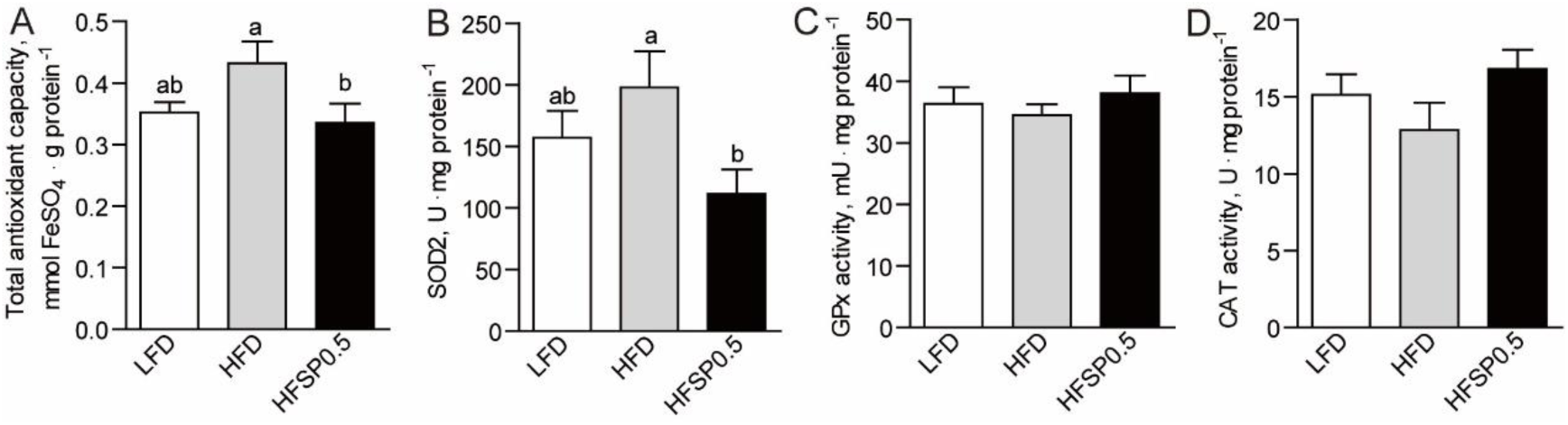
Propionate inhibits intestinal total antioxidant capacity in the context of high fat diet. (A) Intestinal total antioxidant capability in 1-month-old zebrafish fed LFD, HFD or HFSP0.5 diet for 2 wks. (B-D) Intestinal antioxidant enzymes in 1-month-old zebrafish fed LFD, HFD or HFSP0.5 diet for 2 wks, including (B) SOD2, (C) GPx and (D) CAT. Values are means ± SEMs (*n*=5 or 6 biological replicates). Means without a common letter are significantly different, *P*<0.05. SOD2, superoxide dismutase 2; GPx, glutathione peroxidase; CAT, catalase.

### Propionate induces SOD2 propionylation at 132 lysine site

Contrary to lower SOD2 activity, the protein level of intestinal SOD2 in zebrafish fed HFSP0.5 diet was identical to that in zebrafish fed HFD (Fig. 4A and 4B). Since propionate serves as precursor for generation of propionyl-CoA (Schonfeld & Wojtczak, 2016), it seems reasonable to propose that the difference in SOD2 activity between the HFSP0.5 group and the HFD group may involve posttranslational propionylation of SOD2.

**Fig. 4.**
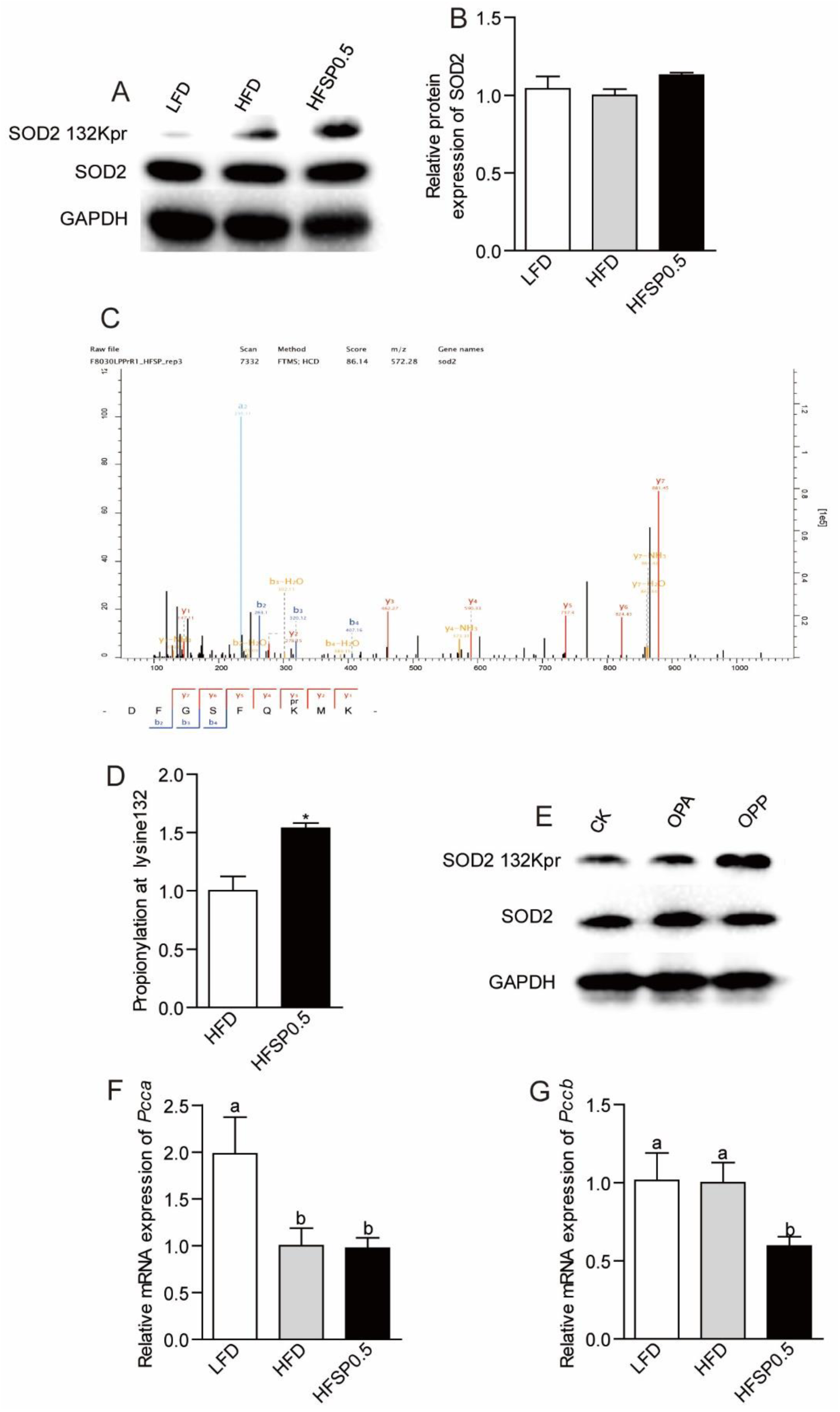
Propionate contributes to SOD2 propionylation at 132 lysine site in the context of high fat diet. (A) A representative western blotting showing patterns of intestinal SOD2 expression and SOD2 propionylation at the 132 lysine site. (B) Quantification of intestinal SOD2 protein level in zebrafish fed LFD, HFD or HFSP0.5 diet for 2 wks. (C) Tandem mass spectrometry from SOD2 demonstrates propionylated lysine 132 *in vivo*. (D) Quantification of intestinal SOD2 propionylation at the 132 lysine site in zebrafish fed HFD or HFSP0.5 diet for 2 wks. (E) A representative western blotting showing patterns of SOD2 expression and SOD2 propionylation at the 132 lysine site in ZF4 cells treated with OPA or OPP. (F-G) The mRNA expression of genes encoding subunits of intestinal PCC, an enzyme catalyzing the carboxylation of propionyl-CoA, in zebrafish fed HFD or HFSP0.5 diet for 2 wks. Values are means ± SEMs, for B and D *n*=2 or 3 biological replicates; for F and G *n*=6 biological replicates. Means without a common letter are significantly different, *P*<0.05. \**P*<0.05, \*\**P*<0.01. OPA, mixture of 150 μM oleic acid and 50 μM palmitic acid; OPP, mixture of 150 μM oleic acid, 50 μM palmitic acid and 50 mM sodium propionate; PCC, propionyl-CoA carboxylase.

Global lysine propionylation performed via HPLC-MS/MS-based proteomics technology showed that SOD2 was the only propionylated antioxidant enzyme in mitochondria (Supplementary Fig. 2). The results showed that SOD2 was propionylated at the 132 lysine site (K132) (Fig. 4A, 4C and 4D). The propionylation of SOD2 K132 was enhanced by exposure of OPP in ZF4 cells (Fig. 4E). Compared with zebrafish fed LFD, zebrafish fed HFD and HFSP0.5 diets showed a lower mRNA level of the PCCa subunit (Fig. 4F). Moreover, zebrafish fed HFSP0.5 diet showed a lower mRNA level of the PCCb subunit compared with zebrafish fed LFD and HFD (Fig. 4G), which indicated that propionate metabolism in the intestine may be disturbed by HFD and HFSP0.5 diets.

### SOD2 propionylation at 132 lysine site accounts for cellular ROS increase

To determine whether propionylation at K132 compromises the activity of SOD2, we generated plasmid expressing mutated zebrafish SOD2 in which K132 was substituted by arginine (R, conserves the positive charge) or glutamine (Q, mimics lysine propionylation) (K132R/Q), and transfected the plasmid into ZF4 cells under SOD2 knockdown state. *Si*RNA (mixture of *Sod2*-1 and *Sod2*-3) targeting SOD2 was used to reduce its expression in ZF4 cells and scrambled *si*RNA was used as a negative control (Supplementary Fig. 3A, Supplementary Table 3). Results showed that overexpression of WT SOD2 and SOD2 K132R/Q compensated the protein level of SOD2 (Fig. 5A). Compared with the cells transfected with WT SOD2, ZF4 cells transfected with SOD K132R mutant showed similar SOD2 activity, ROS level and cell viability (Fig. 5B-5D), while cells transfected with SOD2 K132Q mutant displayed decreased SOD2 activity (Fig. 5B), increased ROS level (Fig. 5C) and lower cell viability (Fig. 5D). These results indicated that propionylation in K132 compromises the enzymic activity of SOD2, leading to enhanced ROS. Moreover, overexpression of SOD2 K132R in ZF4 cells prior to OPP treatment maintained SOD2 activity (Fig. 5E) and prevented cellular ROS elevation (Fig. 5F) when compared to overexpression of WT SOD2, supporting that K132 propionylation induced reduction of SOD2 activity was the main cause of enhanced ROS under high lipid plus propionate conditions. Collectively, these results indicated that propionylation of SOD2 at K132 inhibits SOD2 activity and accounts for cellular ROS accumulation.

**Fig. 5.**
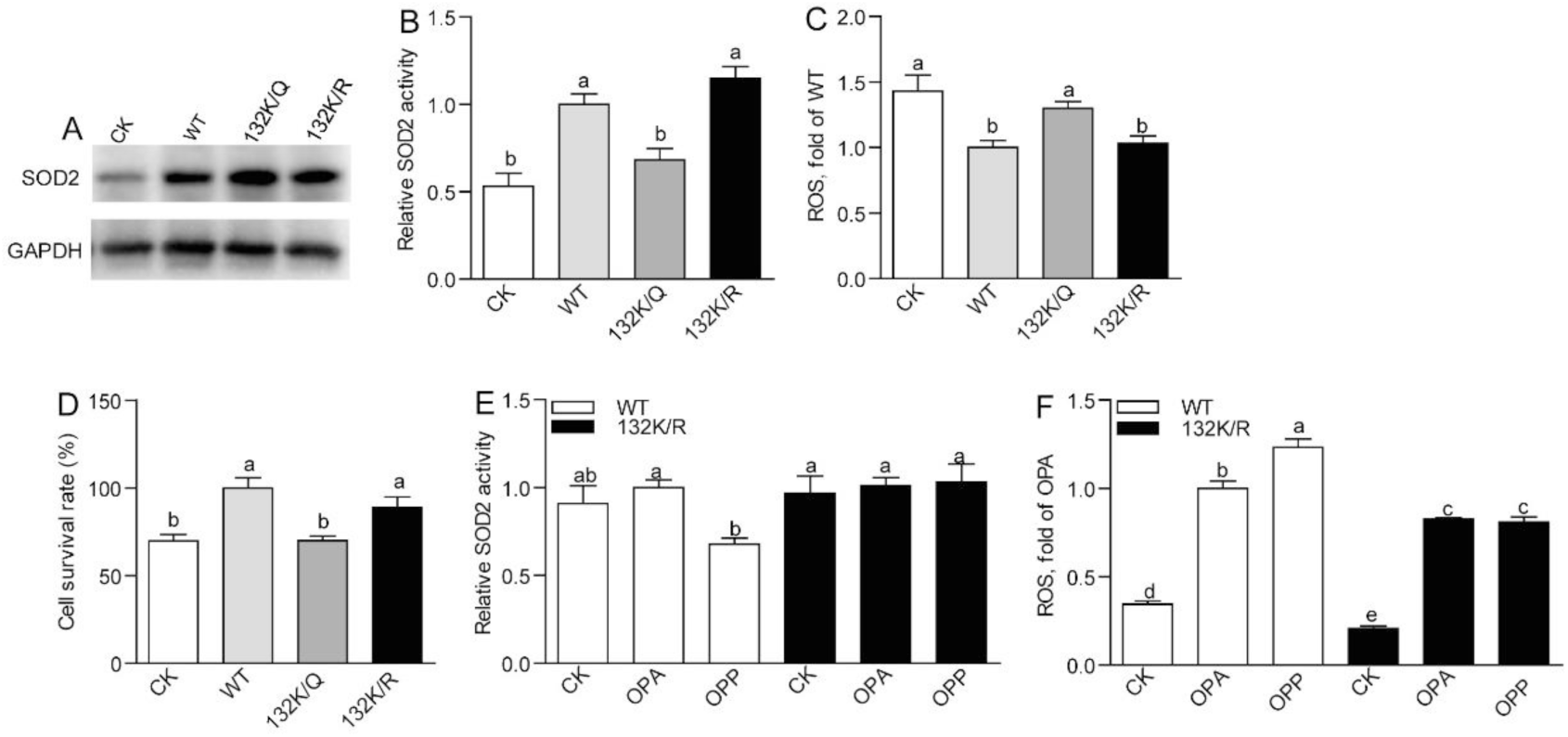
SOD2 propionylation at 132 lysine site accounts for cellular ROS increase. (A) A representative western blotting showing that overexpression of WT SOD2 and SOD2 K132R/Q compensated SOD2 level in ZF4 cells. (B) SOD2 activity, (C) ROS level and (D) cell survival rate in ZF4 cells transfecting with WT SOD2 or SOD K132R/Q mutants. (E) SOD2 activity and (F) ROS level in ZF4 cells treated with OPA or OPP, which were transfected with WT SOD2 and SOD2 K132R in advance. Values are means ± SEMs (*n*=4∼8 biological replicates). Means without a common letter are significantly different, *P*<0.05. OPA, mixture of 150 μM oleic acid and 50 μM palmitic acid; OPP, mixture of 150 μM oleic acid, 50 μM palmitic acid and 50 mM sodium propionate.

### Inhibition of SIRT3 promotes SOD2 propionylation

Recent studies identified that the sirtuin family of deacetylases have depropionylation activity (27 Bheda et al., 2011). To identify which sirtuin is involved in the regulation of SOD2 K132 propionylation, we evaluated the expression of sirtuins in zebrafish intestine. Results showed that the expression of intestinal sirtuin 3 (SIRT3) was reduced in zebrafish fed HFSP0.5 diet when compared with those fed HFD (Fig. 6A-6C). We next examined whether SIRT3 could interact with SOD2 in physiological conditions via immunoprecipitation with intestine lysate. Results showed that SIRT3 could be immunoprecipitated with SOD2 antibody (Fig. 6D). Moreover, exposure of OPP reduced mRNA expression of *Sirt3* in ZF4 cells (Fig. 6E-6G). To identify whether SIRT3 reduction is associated with propionylation of intestinal SOD2 at the K132 site, we knocked down *Sirt3* with *si*RNA (mixture of *Sirt3*-1, *Sirt3*-2 and *Sirt3*-3) in ZF4 cells (Supplementary Fig. 3B, Supplementary Table 3) and detected propionylation of SOD2 at the K132 site via western blot. Results showed that the knockdown of *Sirt3* increased propionylation of SOD2 at K132 (Fig. 6H). In agreement with increased propionylation of SOD2 at K132 in *Sirt3* KD ZF4 cells, the activity of SOD2 (Fig. 6I) and cell viability (Fig. 6J) were significantly reduced. Together, these results indicated that SIRT3 can directly interact with SOD2 and plays an important role in regulating SOD2 activity via modulating propionylation at K132.

**Fig. 6.**
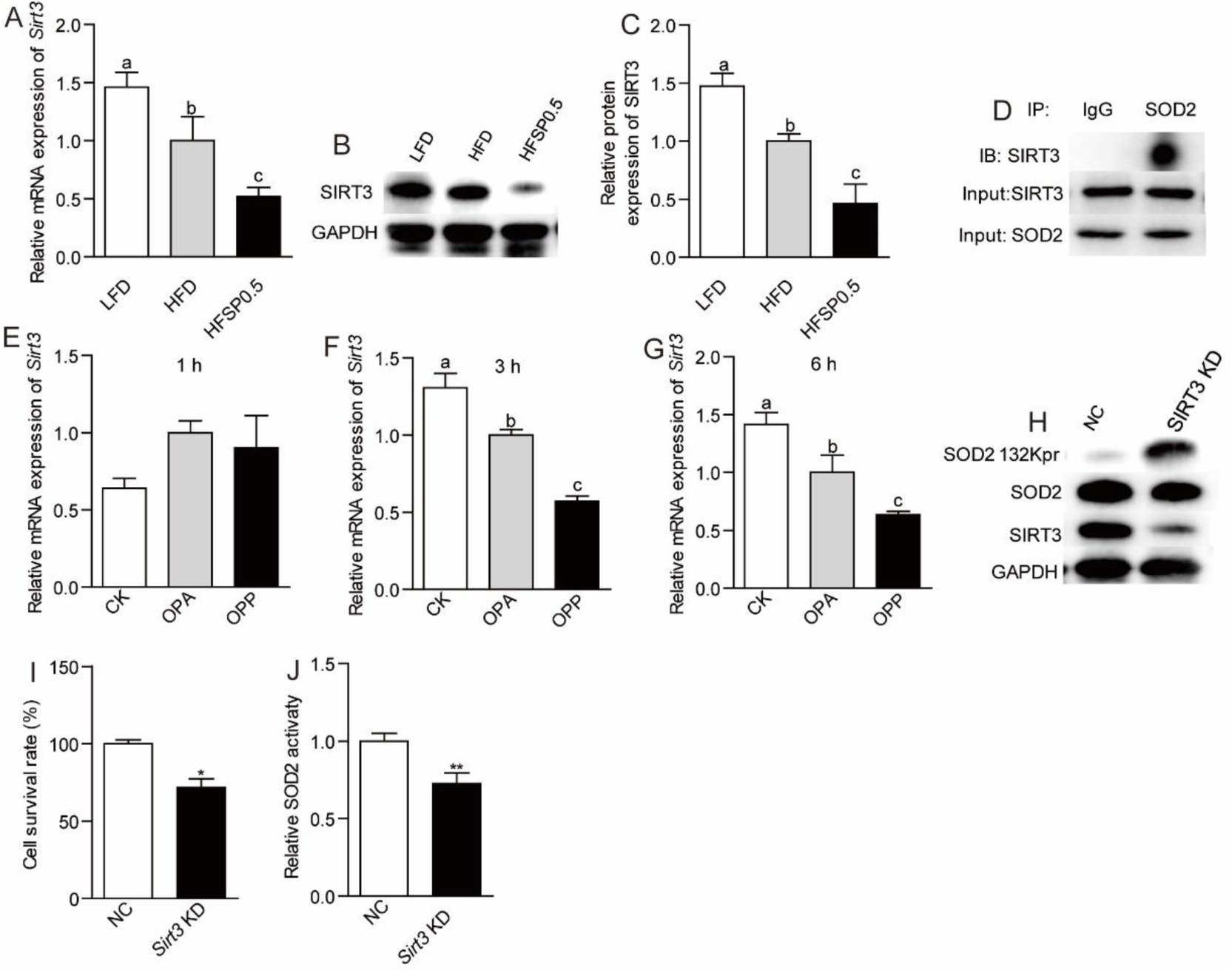
Inhibition of SIRT3 promotes SOD2 propionylation. (A) Intestinal mRNA expression of *Sirt3* in zebrafish fed LFD, HFD or HFSP0.5 diet for 2 wks. (B) A representative western blotting showing expression pattern of intestinal SIRT3. (C) Quantification of intestinal SIRT3 protein level in zebrafish fed LFD, HFD or HFSP0.5 diet for 2 wks. (D) Intestinal SOD2 was immunopurified from intestine lysates with anti-SOD2 antibody, followed by western blotting with anti-SIRT3 antibody. (E-G) The mRNA expression of *Sirt3* in ZF4 cells treated with OPA or OPP in a time-dependent manner. (H) A representative western blotting showing the propionylation of SOD2 at the 132 lysine site in ZF4 cells upon *Sirt3* knockdown. (I) Cell survival rate and (J) SOD2 activity in ZF4 cells upon *Sirt3* knockdown. Values are means ± SEMs, for A *n*=6 biological replicates; for C *n*=3 biological replicates; for E-G *n*=3 or 4 biological replicates; for I and G *n*=6 or 8 biological replicates. Means without a common letter are significantly different, *P*<0.05. \**P*<0.05, \*\**P*<0.01. OPA, mixture of 150 μM oleic acid and 50 μM palmitic acid; OPP, mixture of 150 μM oleic acid, 50 μM palmitic acid and 50 mM sodium propionate.

### Gut microbiota contributes to ROS elevation independent of SIRT3

To determine whether intestinal microbiota is required for the negative effect of propionate in the context of high fat diet, we fed germ free (GF) zebrafish LFD, HFD and HFSP0.5 diet and detected the expression of SIRT3. Results showed that both SIRT3 mRNA level (Fig. 7A) and protein level (Fig. 7B and 7C) in GF zebrafish fed HFSP0.5 diet were significantly lower than those fed HFD. These results indicated that HFSP0.5 diet could directly reduce SIRT3 expression independent of gut microbiota. Consistent with the compromised SIRT3 expression, the propionylation of SOD2 at K132 was enhanced in GF zebrafish fed HFSP0.5 diet (Fig. 7B). Accordingly, SOD2 activity was reduced in GF zebrafish fed HFSP0.5 diet compared with their counterparts fed HFD (Fig. 7D), and ROS was enhanced (Fig. 7E). These results indicated that HFSP0.5 diet could directly induce oxidative stress via SIRT3 inhibition. Although moderately induced in zebrafish fed HFSP0.5 diet compared to those fed HFD, both caspase-9 and caspase-6 activity were significantly higher than in zebrafish fed LFD, and caspase-3 activity was significantly induced in GF zebrafish fed HFSP0.5 diet (Fig. 7F-7H). These results suggest that HFSP0.5 diet activates a mitochondrial death pathway independent of gut microbiota.

**Fig. 7.**
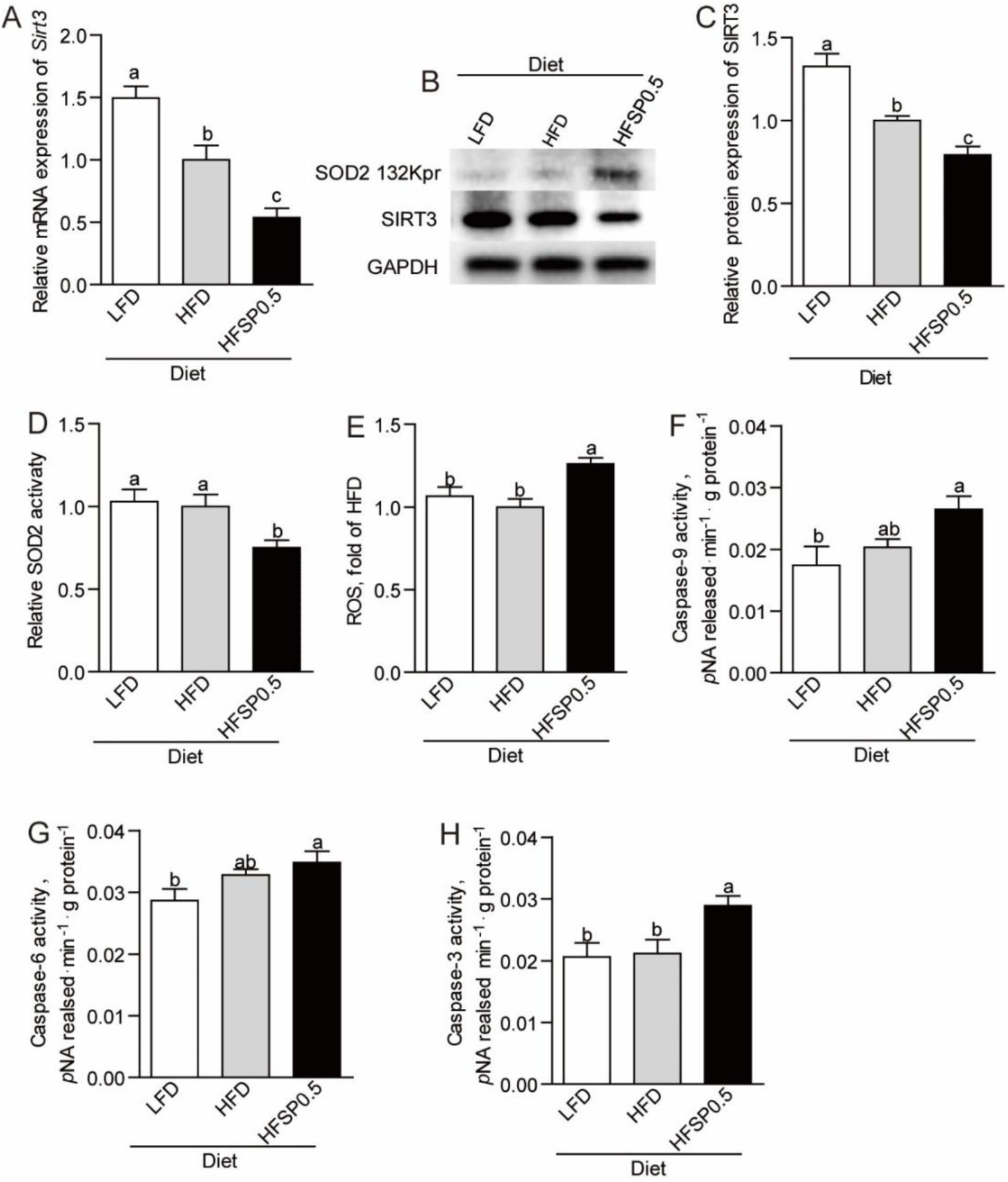
High fat diet supplemented with 0.5% propionate directly inhibits SIRT3 expression. (A) The mRNA expression of *Sirt3* in germ-free (GF) zebrafish fed sterile LFD, HFD or HFSP0.5 diet. (B) A representative western blotting showing patterns of SIRT3 expression and SOD2 propionylation at the 132 lysine site in GF zebrafish fed sterile LFD, HFD or HFSP0.5 diet. (C) Quantification of SIRT3 protein level in GF zebrafish fed sterile LFD, HFD or HFSP0.5 diet. (D) SOD2 activity and (E) ROS level in GF zebrafish fed sterile LFD, HFD or HFSP0.5 diet. The activity of (F) caspase-9, (G) caspase-6 and (H) caspase-3 in GF zebrafish fed sterile LFD, HFD or HFSP0.5 diet. Values are means ± SEMs, for A *n*=4∼6 biological replicates; for C *n*=3 biological replicates; for D-H *n*=4∼8 biological replicates. Means without a common letter are significantly different, *P*<0.05.

To further investigate the role of microbiota, we transferred gut microbiota from zebrafish fed LFD, HFD and HFSP0.5 diet into GF zebrafish and detected the expression of SIRT3. Results showed that gut microbiota from zebrafish fed HFSP0.5 induced moderate increase of SIRT3 expression (Fig. 8A-8C), which was different from the effect of HFSP0.5 diet. These results indicated that the role of gut microbiota may be independent of SIRT3. Gut microbiota from zebrafish fed HFSP0.5 diet induced significant elevation of ROS (Fig. 8D). Meanwhile, the activity of caspase-9, caspase-6 and caspase-3 was significantly higher in GF zebrafish colonized with HFSP0.5-microbiota compared with HFD-microbiota colonized counterparts (Fig. 8E-8G). These results indicated that gut microbiota induced by HFSP0.5 diet could activate the mitochondrial death pathway and induce ROS independent of modulation of SIRT3 expression.

**Fig. 8.**
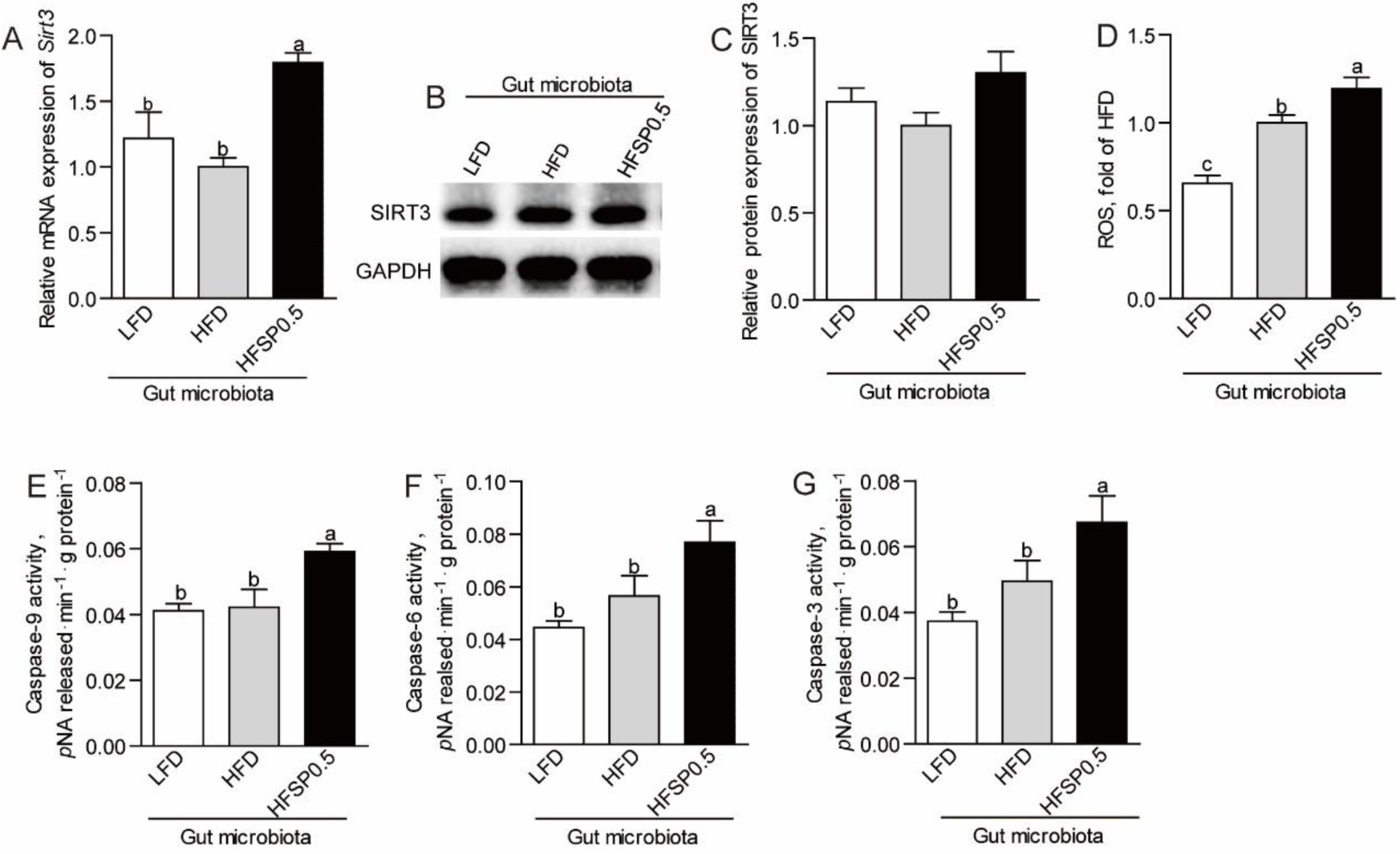
Gut microbiota indirectly activate mitochondrial death pathway. (A) The mRNA expression of *Sirt3* in germ-free (GF) zebrafish transferred with gut microbiota from 1-month-old zebrafish fed LFD, HFD or HFSP0.5 diet. (B) A representative western blotting showing expression patterns of SIRT3 in GF zebrafish transferred with gut microbiota from 1-month-old zebrafish fed LFD, HFD or HFSP0.5 diet. (C) Quantification of SIRT3 protein level in GF zebrafish transferred with gut microbiota from 1-month-old zebrafish fed LFD, HFD or HFSP0.5 diet. (D) ROS level in GF zebrafish colonized with gut microbiota from 1-month-old zebrafish fed LFD, HFD or HFSP0.5 diet. The activity of (E) caspase-9, (F) caspase-6 and (G) caspase-3 in GF zebrafish colonized with gut microbiota from 1-month-old zebrafish fed LFD, HFD or HFSP0.5 diet. Values are means ± SEMs, for A *n*=4 or 5 biological replicates; for C-H *n*=4∼8 biological replicates. Means without a common letter are significantly different, *P*<0.05.

### Alteration of gut microbiota is partly linked to intestinal oxidative stress induced by propionate

Considering the effect associated with the microbiota, we next investigated the mechanism underlying the microbiota alteration. We observed that disturbed luminal redox state was also observed in zebrafish fed HFSP0.5 diet, as evidenced by ROS accumulation in gut content (Fig. 9A). Similarly, ROS level in the medium of ZF4 cells treated with OPP for 24 h was significantly higher than that in cells treated by OPA (Fig. 9B). We analyzed the composition of gut microbiota via 16S *r*RNA gene sequencing, and found that Proteobacteria and *Plesiomonas* were significantly enriched (Fig. 9C and 9D, Tables 1 and 2) in zebrafish fed HFSP0.5 diet compared with those fed HFD (Fig. 9C and 9D, Tables 1 and 2), though total bacterial counts were similar between these two groups (Fig. 9E). The relative abundance of Firmicutes was significantly lower in the HFSP0.5 group when compared with the HFD group (Fig. 9C and Table 1). The relative abundance of Fusobacteria and *Cetobacterium* showed the tendency of decline (Fig. 9C and 9D, Tables 1 and 2).

**Table 1.** The predominant gut bacterial phylum in zebrafish fed on a LFD, HFD, HFSP0.5 diet for four weeks based on V3–V4 sequences.

| Phylum (%) | LFD | HFD | HFSP0.5 |
| --- | --- | --- | --- |
| Proteobacteria | 64.06±5.94b | 73.85±3.10b | 89.70±1.84a |
| Fusobacteria | 14.48±3.82a | 4.65±1.87b | 1.26±0.49b |
| Firmicutes | 17.33±3.52a | 15.90±2.70a | 6.82±1.30b |
| Bacteroidetes | 0.61±0.25 | 1.3±0.62 | 0.62±0.25 |
| Actinobacteria | 2.52±0.63 | 2.32±0.47 | 1.02±0.42 |

Values are expressed as the mean ± SEM, n = 6. Means marked with different letters represent statistically significant results ( $P < 0.05$ ), whereas the same letter correspond to results that show no statistically significant differences.

**Table 2.** The predominant gut bacterial genus in zebrafish fed on a LFD, HFD, HFSP0.5 diet for four weeks based on V3–V4 sequences.

| Genus (%) | LFD | HFD | HFSP0.5 |
| --- | --- | --- | --- |
| Plesiomonas | 17.55±6.92b | 18.28±6.91b | 51.41±7.24a |
| Aeromonas | 9.59±2.37 | 7.60±2.46 | 23.51±2.60 |
| Cetobacterium | 14.27±3.78a | 4.60±4.08b | 1.18±4.15b |
| Lactobacillus | 11.52±2.72a | 3.24±3.27b | 0.62±3.67b |
| Hyphomicrobium | 1.26±0.29b | 4.02±0.27a | 0.38±0.39b |
| Acinetobacter | 1.57±0.86ab | 2.63±0.37a | 0.75±0.10b |
| Chitinibacter | 4.29±4.00 | 0.52±4.01 | 0.06±0.08 |

Values are expressed as the mean ± SEM, n = 6. Means marked with different letters represent statistically significant results ( $P < 0.05$ ), whereas the same letter correspond to results that show no statistically significant differences.

**Fig. 9.**
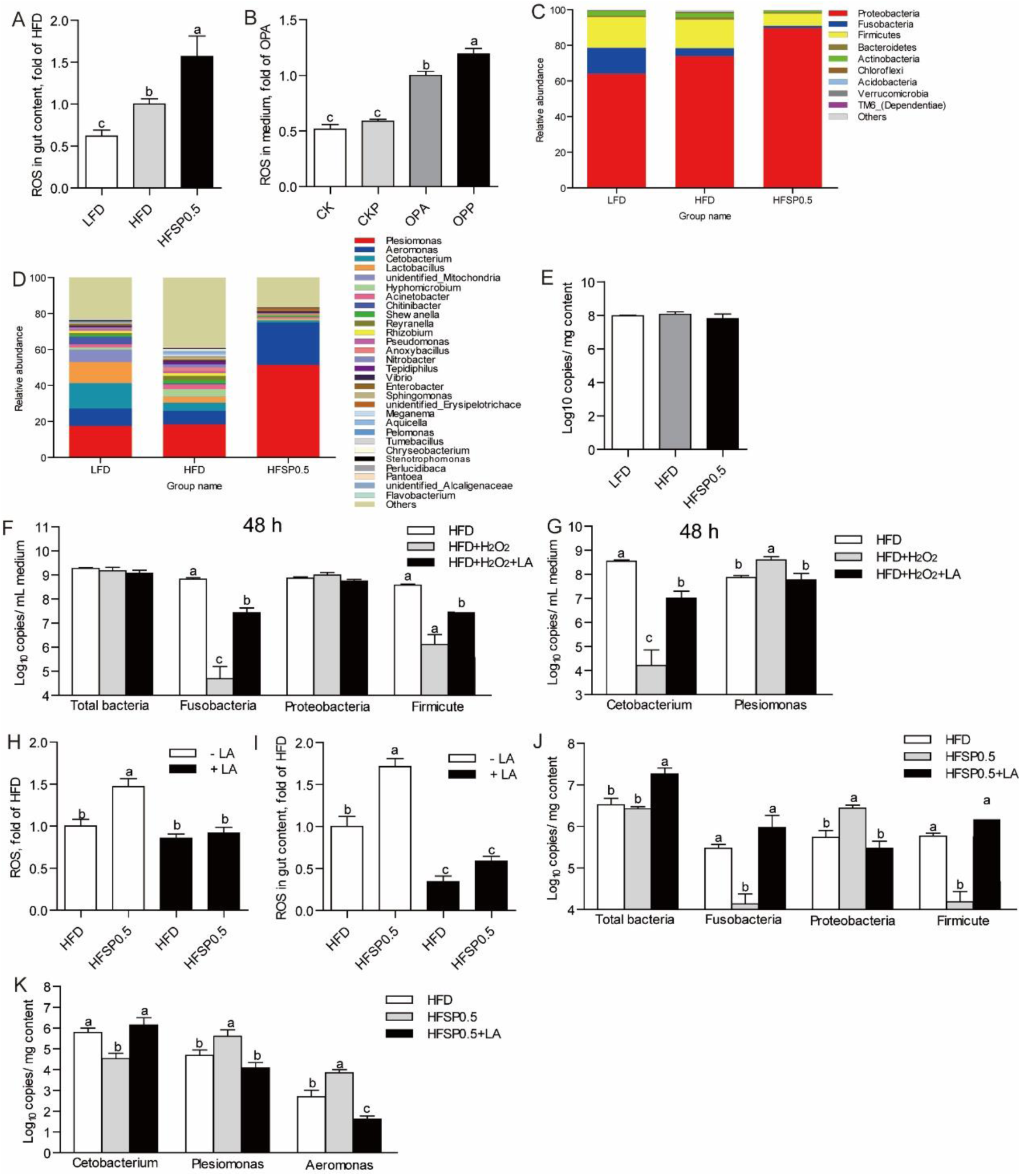
Alteration of gut microbiota is partially linked to intestinal oxidative stress induced by propionate. (A) ROS level in gut content collected from zebrafish fed LFD, HFD or HFSP0.5 diet. (B) ROS level in the medium of ZF4 cells treated with OPA or OPP for 24 h. (C) The composition of gut microbiota at phylum level in 1-month-old zebrafish fed LFD, HFD or HFSP0.5 diet. (D) The composition of gut microbiota at genus level in 1-month-old zebrafish fed LFD, HFD or HFSP0.5 diet. (E) The number of total bacteria (Log10 16S *r*RNA gene copies/mg gut content) in gut content collected from 1-month-old zebrafish fed LFD, HFD or HFSP0.5 diet. (F)The number of total bacteria (Log10 16S *r*RNA gene copies/mL medium), Fusobacteria, Proteobacteria and Firmicutes after incubation in GAM with or without H_2_O_2_ for 48 h. (G) The numbers of *Cetobacterium* and *Plesiomonas* after incubation in GAM with or without H_2_O_2_ for 48 h. (H) ROS level in intestine collected from zebrafish fed HFD or HFSP0.5 diet supplemented with LA. (I) ROS level in gut content collected from zebrafish fed HFD or HFSP0.5 diet supplemented with LA. (J) The number of total bacteria (Log10 16S *r*RNA gene copies/mg gut content), Fusobacteria, Proteobacteria and Firmicutes in gut content collected from 1-month-old zebrafish fed HFSP0.5 diet supplemented with or without LA. (K) The numbers of *Cetobacterium, Plesiomonas* and *Aeromonas* in gut content collected from 1-month-old zebrafish fed HFSP0.5 diet supplemented with or without LA. Values are means ± SEMs (*n*=4∼6 biological replicates). Means without a common letter are significantly different, *P*<0.05.

To determine the link between disturbed luminal redox state induced by HFSP0.5 diet and the composition of gut microbiota, we evaluated the effect of oxidative stress on gut microbiota *in vitro* by culturing gut microbiota isolated from zebrafish fed HFD in H_2_O_2_-supplemented GAM. Results showed that when incubated in GAM containing 2 mmol/L H_2_O_2_ for 48 h, the numbers of Fusobacteria, Firmicutes and *Cetobacterium* were dramatically decreased, while Proteobacteria showed a moderate increase and the number of *Plesiomonas* was significantly elevated (Fig. 9F and 9G). Meanwhile, 0.5 mg/mL lipoic acid (LA), a universal antioxidant, maintained the numbers of Fusobacteria, Firmicutes and *Cetobacterium* similar to those cultured in GAM without H_2_O_2_ and restricted the growth of *Plesiomonas* (Fig. 9F and 9G). These results indicated that disturbed redox state may contribute to the compositional change of the intestinal microbiota in zebrafish.

To further validate the effects of oxidative stress induced by HFSP0.5 diet on gut microbiota, we performed *q*PCR to identify the numbers of Proteobacteria and *Plesiomonas* in gut content collected from zebrafish fed HFSP0.5 diet with or without LA. Results showed that ROS accumulation in intestine and gut content collected from zebrafish fed HFSP0.5 diet was significantly alleviated by LA supplementation (Fig. 9H and 9I). Accordingly, the numbers of Proteobacteria and *Plesiomonas* in gut content were significantly decreased by LA compared with the control (Fig. 9J and 9K). Together, these results indicated that oxidative stress induced by propionate in the context of HFD has the potential to switch the composition of gut microbiota by elevating the abundance of Proteobacteria, which in turn further activate the mitochondrial death pathway and exacerbate oxidative stress.

## Discussion

Emerging evidence suggests that propionate can be a dietary factor to ameliorate diet-induced obesity (Lin et al., 2012; Lu et al., 2016; den Besten et al., 2015) and reduce liver lipogenesis (Wright et al., 1990; Weitkunat et al., 2016). In this study, the anti-obesity effect of propionate was also observed in zebrafish fed HFSP0.5 diet, as evidenced by lower body weight gain and hepatic lipid accumulation. However, oxidative stress in the intestine happened together with the anti-obesity effect of propionate, as shown by elevated ROS, MDA and PC. Oxidative stress is caused by ROS accumulation due to the imbalance of ROS production and removal, resulting in mitochondrial dysfunction (Wei et al., 1998; Duchen, 2004; Pieczenik & Neustadt, 2007). SOD2 is the primary mitochondrial enzyme for ROS clearance (Spitz & Oberley, 1989; Zelko et al., 2002). Our study suggests that propionate impairs intestinal anti-oxidant capability and SOD2 activity in the context of HFD.

So far, the enzyme activity of SOD2 has been shown to be regulated by an ubiquitous post translational modification (PTM), acetylation (Chen et al., 2011; Assiri et al., 2016; Liu et al., 2017). Among all lysine residues of SOD2 in mammals, lysine sites 53, 68, 89, 122 and 130 have been shown to be acetylated (Qiu et al., 2010; Tao et al., 2010; Lu et al., 2015; Zhang et al., 2016). According to the alignment of amino acid sequences of SOD2 among zebrafish (EMBL no. AY195857), mice (AK002534), rats (BC070913) and humans (M36693), the 132 lysine of zebrafish SOD2 is aligned to the 130 lysine of mouse, rat and human SOD2 (Lin et al., 2009). Moreover, SIRT3-mediated deacetylation of SOD2 (K130) can prevent ROS accumulation (Zhang et al., 2016). Our results identify intestinal SOD2 lysine 132 as the propionylated lysine site induced by 2-week propionate feeding under HFD. In addition, SOD2 activity could be modulated by propionylation of K132, which was validated by K132Q and K132R mutants that demonstrated decreased activity when lysine 132 was replaced by Gln to mimic lysine propionylation, as well as decreased propionate-induced ROS level when lysine 132 was replaced by Arg to mimic lysine depropionylation. Thus, lysine 132 in zebrafish SOD2 is a key residue which is important for regulation of SOD2 activity.

Apart from SOD2, lysine propionylation was also observed in other proteins (Supplementary Fig. 2) based on the global lysine propionylation analysis (Supplementary Fig. 2). The first global survey of lysine propionylation has been reported in Cyanobacteria (Yang et al., 2019); however, there has been no report on global propionylome in animals. The bioinformatics results showed that proteins involved in oxidative phosphorylation (OXPHOS) and ATP synthesis were enriched among the propionylated proteins in HFSP0.5-zebrafish intestine. Besides, proteins associated with the KEEG pathway of the citrate cycle (TCA cycle) were enriched for lysine propionylation. Among these proteins in the TCA cycle, propionylated malate dehydrogenase 2 (MDH2) and citrate synthase (CS) were found to have a complex association with other mitochondrial proteins. This may indicate the potential regulatory role of lysine propionylation in mitochondrial metabolism via modulating functions of MDH2 and CS. SOD2 was the only propionylated antioxidant enzyme in mitochondria, which was consistent with our results that blocking SOD2 propionylation at K132 prevented ROS elevation in OPP-treated ZF4 cells. Nevertheless, our results showed that global lysine propionylation may potentially modulate mitochondrial energy metabolism, which deserves further investigation.

SIRT3 is a mitochondrially localized deacetylase (Michishita et al., 2005) which has been reported as a central regulator of mitochondrial ROS production and is required for protection from oxidative damage (Bause & Haigis, 2013). Cells lacking SIRT3 are susceptible to oxidative stress (Qiu et al., 2010; Wang et al., 2014). We show that propionate-induced SOD2 K132 propionylation is accompanied by depressed SIRT3 expression and that SOD2 interacts with intestinal SIRT3. The sirtuin family of deacetylases has been reported to have depropionylation activity. For instance, the propionyl-lysine modification introduced by bacterial Gcn-5-related N-acetyltransferase enzymes can be removed by bacterial and human sirtuins (Garrity et al., 2007). Moreover, the absence of SIRT3 leads to a higher propionylated lysine level in mouse lenses (Nahomi et al., 2020). In this study, ZF4 cells with depressed SIRT3 exhibited an elevated propionylation level in SOD2 132 lysine, as well as impaired SOD2 activity. These results suggest a PTM mechanism involving propionylation for SIRT3 modulation of SOD2 that is independent of SOD2 expression. Several studies suggest that SIRT3 expression is dynamically regulated by nutrition, such as caffeine and diet restriction (Zhang et al., 2015; Yu et al., 2018). Peroxisome proliferator-activated receptor γ coactivator 1 α (PGC1a) is one of few known regulators of SIRT3, which can activate SIRT3 expression by binding the ERR binding element in the promoter region (Kong et al., 2010). The reported transcriptional repressors including poly (ADP-ribose) polymerase 1 (PARP1) and transcriptional cofactor receptor-interacting protein 140 (RIP140), both of which contribute to oxidative stress and mitochondrial dysfunction (Yoon & Kim, 2016; Kim et al., 2020). In this study, the expression of *Pgc1α* and *Err* in zebrafish intestine and ZF4 cells showed no significant alteration in response to HFSP0.5 diet and OPP treatment (Supplementary Fig. 4), suggesting that the reduction of SIRT3 expression might be mediated by its transcriptional repressors.

Germ-free zebrafish is a convenient tool to investigate the contribution of dietary factors or gut microbiota on host health and disease. Our studies demonstrate an inhibitory mode to SIRT3 in the GF zebrafish model, which is independent of gut microbiota. Thus, oxidative stress resulting from SOD2 propionylation is independent of gut microbiota. On the other hand, gut microbiota induced by HFSP0.5 diet increased ROS accumulation. Microbiota targets mitochondria to regulate interaction with the host, and the mitochondrial production of ROS is often targeted by pathogenic bacteria (Saint-Georges-Chaumet & Edeas, 2016). Pathogenic bacteria release various pathogen-associated molecular patterns (PAMPs), including lipopolysaccharides (LPS), flagellin, lipoteichoic acid, lipoprotein or other toxins, which can be recognized by the pattern recognition receptor (PRR) system in the host cell surface and further induce mitochondrial ROS production (Saint-Georges-Chaumet & Edeas, 2016; Emre & Nubel, 2010). Therefore, the alteration in the microbiota structure might promote ROS production due to differential microbe-associated molecular patterns (MAMPs)-PRR signaling.

Gut microbiota tends to be influenced by dietary macronutrients or the microenvironment in the lumen (Conlon & Bird, 2016; Hevia et al., 2015). Given that the contents of fat, carbohydrates and protein were consistent between the HFD and HFSP0.5 diets, the influence of dietary macronutrients can have been negligible. Thus, changes of the gut microenvironment in zebrafish fed HFSP0.5 diet may have driven the alteration of gut microbiota. In the present study, both intestinal oxidative stress and abnormal luminal redox state occurred in zebrafish fed HFSP0.5 diet and could be eliminated by LA. The reason for disturbed luminal redox state remained undiscovered, but elevated ROS level in the medium of ZF4 cells treated with OPP showed that at least part of ROS in lumen is released by intestinal cells. Gut microbiota in zebrafish fed HFSP0.5 diet was characterized with enrichment of Proteobacteria and *Plesiomonas*. Since HFSP0.5 diet-associated alteration of gut microbiota can be restored by LA *in vivo*, it is reasonable to propose that luminal redox state is linked to gut microbiota alteration. Besides, the quantitative results for gut microbiota cultured *in vitro* showed that the numbers of Fusobacteria and Firmicutes were reduced by H_2_O_2_, whereas the Proteobacteria population showed a modest increase. This suggested that Proteobacteria are more resilient to oxidative stress than Fusobacteria and Firmicutes, which may explain the enrichment of Proteobacteria in a disturbed redox state. Our results indicated that gut microbiota acted both as a responder and as a secondary inducer of the intestinal oxidative stress, while the original cause was attributed to propionate and HFD.

One intriguing finding in our work is that propionate only induces oxidative damage to the intestine under a high fat background. Considering that propionate-induced oxidative damage is mediated by the modulation of SOD2 lysine 132 propionylation, SIRT3 and propiony-CoA metabolism are important to illuminate the effect of dietary fat on propionate toxicity. SIRT3 expression can be depressed by nutritional stress, such as alcohol and HFD (Ma et al., 2019; Palacios et al., 2009). Mitochondrial protein propionylation increases in response to chronic ethanol ingestion in mice, similar to mitochondrial protein acetylation (Fritz et al., 2013). In this study, we show that intestinal SIRT3 expression in zebrafish fed HFD is lower than in zebrafish fed LFD. Propionyl-CoA carboxylase (PCC) is the essential enzyme catalyzing the carboxylation of propionyl-CoA to methylmalonyl-CoA, ultimately contributes to the succinyl-CoA pool and enters the TCA cycle (Wongkittichote & Chapman, 2017; Xu et al., 2018). Our results show that HFD inhibits PCC expression, which may disrupt the conversion of propionyl-CoA to succinyl-CoA. Deficiency of PCC leads to accumulation of propionyl-CoA (Wongkittichote & Chapman, 2017). The results presented above suggest that the capability of intestine for depropionylation and metabolism of propionyl-CoA are weakened by HFD. Although the contribution of HFD to SOD2 propionylation remains unclear, a high level of dietary fat promotes the potential adverse effects of propionate.

In clinical research, individuals with ASD are four times as likely to experience gastrointestinal (GI) symptoms as healthy controls (McElhanon et al., 2014). An important factor in the pathogenesis of ASD is gut microbiota, such as *Clostridium* spp., and its metabolites, especially propionate (Wang et al., 2014). There is some evidence that gut microbiota (Strati et al., 2017; Finegold et al., 2010), increased intestinal inflammation (Nina & David, 2017) and mitochondrial dysfunction (Frye et al., 2015) may play a role in ASD-associated GI symptoms, but the pathogenesis has not been well defined. The results in our study suggest that propionate-induced propionylation of antioxidant proteins and the resultant intestinal oxidative stress may contribute to GI-related comorbidities in ASD. In another metabolic disorder known as propionic acidemia (PA), the accumulation of propionyl CoA results in mitochondrial dysfunction and oxidative stress (de Keyzer et al., 2009; Gallego-Villar et al., 2013; Gallego-Villar et al., 2016). Meanwhile, fibroblasts from patients with PA show increased protein propionylation, which leads to impaired mitochondrial respiration (Pougovkina, 2016). Although studies suggest the role of protein propionylation and oxidative stress in the pathological mechanism of PA, how protein propionylation manipulates oxidative stress was unmentioned and the identification of propionylated proteins has been lacking. Our results identified propionylated SOD2 as a direct cause of oxidative stress, implying that a similar mechanism may apply to PA pathogenesis.

The results presented above suggest that propionate in the context of high fat diet induces SOD2 propionylation at the 132 lysine site, which compromises the superoxide scavenging function of SOD2 and induces oxidative stress in zebrafish intestine. SIRT3 can directly interact with SOD2 and plays an important role in regulating SOD2 activity via modulating propionylation at K132, and the enhanced SOD2 propionylation in zebrafish fed high fat plus propionate diet was attributed to reduced SIRT3 expression. Although the connection between SIRT3-mediated deacetylation and SOD2 activation has been well demonstrated, this is the first identification of the link between SIRT3-mediated depropionylation and SOD2 activity. Furthermore, our results indicated that the intestinal microbiota associated with high fat plus propionate diet also induces oxidative stress in zebrafish intestine, but in a SIRT3-independent manner, while the oxidative stress contributes to the shaping of the microbiota. Considering propionate is a widely used food and feed preservative, our findings have important implications for the safety of propionate as a food additive, especially in the context of high fat diet.

## Materials and Methods

### Fish husbandry

All of the experimental and animal care procedures were approved by the Feed Research Institute of the Chinese Academy of Agricultural Sciences Animal Care Committee under the auspices of the China Council for Animal Care (Assurance No. 2016-AF-FRI-CAAS-001). One-month-old zebrafish were maintained at the zebrafish facility of the Feed Research Institute of the Chinese Academy of Agricultural Sciences (Beijing, China) and fed with the experimental diets (Supplementary Table 1) twice a day (9:00, 17:00) to apparent satiation each time for 2 weeks. During the feeding period, the rearing temperature was 25-28 °C, the dissolved oxygen was > 6.0 mg/L, the pH was 7.0 - 7.2, the nitrogen content was < 0.50 mg · L^-1^, and the nitrogen content (as NO_2_) was < 0.02 mg · L^-1^. All fish were anesthetized with tricaine methanesulfonate (MS222).

### Examination of intestinal histopathology

The intestines of zebrafish were rinsed with sterilized PBS, fixed in 4% formalin solution, and embedded in paraffin. For histological analysis, the liver sections prepared from the paraffin blocks were stained with hematoxylin and eosin (H&E). Images were obtained under a microscope (Carl Zeiss) at a 200× magnification.

### Intraperitoneal injection of 4-Hydroxy-TEMPO

After 2-wk feeding trial, ten zebrafish from LFD, HFD and HFSP0.5 groups were respectively divided into two groups: 1) control zebrafish injected intraperitonealy (i.p.) with saline (0.9% NaCl); 2) zebrafish treated i.p. with Tempol dissolved in saline (10 mg/kg b.w.) every other day. Intestines were collected at the sixth day for H&E staining.

### Detection of caspase activity

The activities of caspase-3, caspase-6 and caspase-9 were determined using an assay kit (Beyotime Biotechnology, Shanghai, China) according to the manufacturer’s instructions. The optical density of the reaction product was examined at 405 nm. The enzyme activity units were expressed as the rate of *p*-nitroaniline (*p*NA) released from the substrate per gram protein (μmol *p*NA released · min^-1^ · g protein^-1^).

### Mitochondria isolation and mitochondrial reactive oxygen species determination

Intestine was used to perform mitochondria isolation by using a tissue mitochondria isolation kit (Beyotime Biotechnology, Shanghai, China) according to the manufacturer’s instructions. We adjusted mitochondria to the same level according to mitochondria protein level. Mitochondria isolated were then subjected to reactive oxygen species (ROS) assay using fluorometric intracellular ROS kit (Sigma, USA). The fluorescence was acquired with excitation and emission wavelengths of 490 nm and 520 nm, respectively. ROS level was expressed as the fold of the HFD group.

### Detection of oxidative parameters

Lipid peroxidation was determined by the reaction of malondialdehyde (MDA) with thiobarbituric acid to form a colorimetric product by using a lipid peroxidation assay kit according to the manufacturer’s instructions (Sigma, USA). The optical density of the reaction product was examined at 532 nm. Lipid peroxidation was expressed as MDA content per 100 milligram tissue (nmol · 100 mg tissue^-1^). Oxidation of proteins was determined by the formation of stable dinitrophenyl hydrazine adducts derived from protein carbonyl (PC) groups with 2, 4-dinitrophenylhydrazine using protein carbonyl content assay kit according to the manufacturer’s instructions (Sigma, USA). The optical density of the adduct was examined at 375 nm. Oxidation of proteins was expressed as PC content per milligram protein (nmol · mg protein^-1^).

### Evaluation of total antioxidant capacity

Total antioxidant capacity (T-AOC) was measured by the production of blue Fe^2+^-TPTZ resulting from the reduction of Fe^3+^ TPTZ complex in acidic conditions. The optical density was measured at 593 nm. T-AOC was defined as the production of FeSO_4_ per gram protein (mmol FeSO_4_ · g protein^-1^).

### Evaluation of antioxidant enzyme activity

The activity of SOD2, glutathione peroxidase (GPx) and catalase (CAT) were detected using a CuZn/Mn-SOD activity kit, a cellular GPx assay kit and a catalase assay kit (Beyotime Biotechnology, Shanghai, China), respectively, according to the manufacturer’s instructions. SOD2 activity was measured as the inhibition of water soluble tetrazol salt (WST-8) reduction in a xanthine-xanthine oxidase system. The SOD2 activity in intestine was expressed as U · mg protein^-1^. Relative SOD2 activity in ZF4 cell and zebrafish larva was expressed as fold of indicated group. The activity of GPx was expressed as mU · mg protein^-1^. The activity of CAT was expressed as U · mg protein^-1^.

### Gut microbiota analysis

The 16s V3 – V4 region was amplified using the primers U341F (5’-CGGCAACGAGCGCAACCC-3’) and U806 (5’-CCATTGTAGCACGTGTGTAGCC-3’). The 16S ribosomal RNA gene sequencing was performed by Novogene Bioinformatics Technology Co. Ltd (Beijing, China) using the Illumina HiSeq platform. Then the raw pair-end readings were subjected to a quality-control procedure using the UPARSE-operational taxonomic unit (OTU) algorithm. The qualified reads were clustered to generate OTUs at the 97% similarity level using the USEARCH sequence analysis tool. A representative sequence of each OTU was assigned to a taxonomic level in the Ribosomal Database Project (RDP) database using the RDP classifier. Principal component analysis and heat-map analysis were performed by using R 3.1.0.

### GF-zebrafish generation and treatment

GF-zebrafish were derived from normal zebrafish and reared following established protocols (Rawls et al., 2006). We formulated microparticulate diets, namely LFD, HFD, and HFSP0.5 diets, for zebrafish larvae (Supplementary Table 2). Before feeding, the microparticulate diets were sterilized by irradiation with 20 kGy gamma ray in an atomic energy center (Institute of Food Science and Technology, Chinese Academy of Agricultural Sciences, Beijing, China). Zebrafish larvae hatched from their chorions at 3 days postfertilization (dpf). Each group had six bottles with 20 fish per bottle. At 5 dpf, the yolk was largely absorbed and the GF-zebrafish started feeding. At 11 dpf, whole fish were collected for analysis of caspase activity, SOD2 activity, *q*PCR or western blotting.

### Cell culture

The ZF4 cell line was purchased from American Type Culture Collection (Manassas, VA, USA), and cultured according to established protocols (Driever & Rangini, 1993). The media were obtained from Corning Inc. (New York, NY, USA). Penicillin-Streptomycin solution and bovine insulin were purchased from Sigma (St. Louis, MO, USA). Fetal bovine serum was purchased from Corning Inc. (New York, NY, USA).

### Cell viability analysis

ZF4 cell was first seeded on 96-well plates and incubated for 24 h to sub-confluence. Then ZF4 cell was exposed to fresh medium added with a mixture of 150 μM OA and 50 μM PA (OPA), and a mixture of 150 μM OA, 50 μM PA and 50 mM propionate (OPP). At the end of the exposure period, fresh medium with 10% AlarmaBlue cell viability reagent (Invitrogen, Grand Island, NY, USA) was added. After a 1-h incubation, fluorescence was measured with the SynergyMX Multi-Functional Detector (Biotek, Winooski, VT, USA) at excitation and emission wavelengths of 485 nm and 595 nm, respectively. The ratio of cell viability was calculated using the fluorescence readings of the control and treatments.

### Cell apoptosis analysis

Cell apoptosis detection was performed with Annexin V-fluorescein isothiocyanate (FITC) kits (Sigma). After exposure to OPA or OPP for 24 h, the cells were collected and incubated with Annexin V-FITC and propidium iodide in binding buffer for 10 min in darkness at room temperature. The analysis was conducted by the Guava easyCyte Flow Cytometer (Merck Millipore, Stafford, VA, USA).

### Gene silencing with *si*RNA

Scrambled *si*RNA (negative control), *Sod2* and *Sirt3 si*RNA (Supplementary Table 3) were synthesized by GenePharma Co. Ltd. (Shanghai, China). Cells were first seeded on 6-well plates (Corning) and incubated for 24 h to sub-confluence. Then the cells were transfected with the appropriate *si*RNAs using Lipofectamine RNAiMAX Transfection Reagent (Invitrogen). Efficiency of the *si*RNA was determined by *q*PCR.

### Effects of oxidative stress on gut microbiota *in vitro*

Fresh gut content samples pooled from five zebrafish were put on ice and diluted in 5 mL sterile, ice-cold PBS. Within 30 min of sample collection, bacteria were cultured on Gifu anaerobic medium (GAM); GAM supplemented with 2 mmol/L H_2_O_2_ (GAM+H_2_O_2_); GAM+H_2_O_2_+LA, GAM supplemented with 2 mmol/L H_2_O_2_ and 0.05 % LA. After an incubation period of 48 h at 28 °C, the number of total bacteria or a specific phylotype was quantified by *q*PCR according to Zhang et al (73 Zhang et al., 2019). Primer sets for universal bacteria or specific bacterial groups targeting the 16S *r*RNA gene are listed in Supplementary Table 4. For the gut microbiota cultured in vitro, results were expressed as Log10 copy numbers of bacterial 16S rDNA per mL medium (Log_10_ copies/mL medium).

### Plasmid construction and transfection

The SOD2 was cloned into *p*CDNA3.1. Point (Site) mutations of SOD2 were generated by QuikChange Site-Directed Mutagenesis kit (Stratagene). Both WT SOD2 and SOD2 mutant plasmids were transfected into ZF4 cells using Lipofectamine 3000 Transfection Reagent (Invitrogen).

### Western blotting

Zebrafish intestine or larval zebrafish were homogenized in ice-cold HBSS buffer mixed with 1 mM PMSF and phosphatase inhibitors. Equivalent amounts of total protein were loaded into a 12% SDS-PAGE for electrophoresis and then transferred into a PVDF membrane (Millipore, USA). After blocking nonspecific binding with 5% skimmed milk in TBST, the PVDF membrane was incubated with primary antibodies, i.e., antibodies against GAPDH (Sigma, SAB2708126, 1:2000), SOD2 (Genetex, GTX124294, 1:1000), SIRT3 (Sigma, AV32388, 1:1000) and customized SOD2 k132pro (Jingjie, 1:500). The blots were developed using HRP-conjugated secondary antibodies (GE Health, 1:3000) and the ECL-plus system.

### Total RNA extraction, reverse transcription, and *q*PCR

Total RNA was isolated using Trizol reagent and then reverse transcribed to cDNA. The *q*PCR was performed using SYBR®Green Supermix according to the manufacturer’s instructions (Tiangen, Beijing, China). The results were stored, managed, and analyzed via LightCycler 480 software (Roche, Basel, Switzerland). The *q*PCR primers used are listed in Supplementary Table 4.

### Data analysis

All of the statistical analyses were conducted using GraphPad Prism 5 software (GraphPad Software Inc., San Diego, CA, USA). Results are expressed as the means ± standard errors of the means (SEMs). Comparisons between two groups were analyzed using the Student’s *t*-test, and comparisons between multiple groups were analyzed using one-way ANOVA followed by a Duncan’s test. The statistical significance was set at *P*<0.05.

## Acknowledgments

This work was supported by the National Natural Science Foundation of China (NSFC 31925038, 31972807, 31872584, 31802315) and the National Key R&D Program of China (2018YFD0900400) and the National Natural Science Foundation of China (NSFC 31802315, 31760762, 31872584 and 31702354) funded this work for grants to Dr. Zhigang Zhou, and NIBIO’s STIM China grant (Grant No. 51133) to Dr Jihong Liu Clarke. The authors thank Prof. Nicholas Clarke for linguistic proof reading.

## Author contributions

Z.Z.G. designed the research. D.Q.W and R.C. wrote the paper, and Z.Z.G. gave conceptual advice for the paper. J.L.C. reviewed and helped to revise the manuscript. D.Q.W performed experiments and acquired data. Z.Z. and Y.L. assisted in the *q*PCR, western blot, gut microbiota analysis and *si*RNA knockdown experiments. L.H.L. and H.Q. participated in zebrafish husbandry and sampling. R.C., Y.Y.L. and Z.Z. co-analyzed and discussed the results. All authors read and approved the final manuscript.

