## Supplementary data for "Dietary propionate induces intestinal oxidative stress via inhibition of SIRT3-mediated SOD2 depropionylation"

#### Supplementary Methods

##### Identification of lysine propionylated sites by HPLC-MS/MS

The intestine was ground by liquid nitrogen into cell powder and then transferred to a 5 mL centrifuge tube. After that, four volumes of lysis buffer (8 M urea, 1% protease inhibitor cocktail, 3 μM trichostation, 50 mM nicotinamide) were added to the cell powder, followed by sonication three times on ice using a high intensity ultrasonic processor. The remaining debris was removed by centrifugation at 12,000 g at 4°C for 10 min. Finally, the supernatant was collected and the protein concentration was determined with BCA kit according to the manufacturer's instructions. For digestion, the protein solution was reduced with 5 mM dithiothreitol for 30 min at 56 °C and alkylated with 11 mM iodoacetamide for 15 min at room temperature in darkness. The protein sample was then diluted by adding 100 mM TEAB to urea concentration less than 2M. Finally, trypsin was added at 1:50 trypsin-to-protein mass ratio for the first digestion overnight and 1:100 trypsin-to-protein mass ratio for a second 4 h digestion. To enrich propionylation modified peptides, tryptic peptides dissolved in NETN buffer (100 mM NaCl, 1 mM EDTA, 50 mM Tris-HCl, 0.5% NP-40, pH 8.0) were incubated with pre-washed antibody beads (PTM202, PTM Bio) at 4 °C overnight with gentle shaking. Then the beads were washed four times with NETN buffer and twice with $H_2O$. The bound peptides were eluted from the beads with 0.1% trifluoroacetic acid. Finally, the eluted fractions were combined and vacuum-dried. For HPLC-MS/MS analysis, the resulting peptides were desalted with C18 ZipTips (Millipore) according to the manufacturer's instructions.

The tryptic peptides were dissolved in 0.1% formic acid (solvent A), directly loaded onto a home-made reversed-phase analytical column (15 cm length, 75 μm i.d.). The gradient was comprised of an increase from 6% to 23% solvent B (0.1% formic acid in 98% acetonitrile) over 26 min, 23% to 35% in 8 min and climbing to 80% in 3 min then holding at 80% for the last 3 min, all at a constant flow rate of 400 nL/min on an EASY-nLC 1000 UPLC system. The peptides were subjected to NSI source followed by tandem mass spectrometry (MS/MS) in Q ExactiveTM Plus (Thermo) coupled online to the UPLC. The electrospray voltage applied was 2.0 kV. The m/z scan range was 350 to 1800 for full scan, and intact peptides were detected. The resulting MS/MS data were processed using Maxquant search engine (v.1.5.2.8). Tandem mass spectra were searched against the *Danio rerio* database concatenated with a reverse decoy database. Trypsin/P was specified as cleavage enzyme allowing up to 4 missing cleavages. The mass tolerance for precursor ions was set as 20 ppm in first search and 5 ppm in main search, and the mass tolerance for fragment ions was set as 0.02 Da. Carbamidomethyl on Cys was specified as fixed modification and propionylation modification and oxidation on Met were specified as variable modifications. FDR was adjusted to < 1% and minimum score for modified peptides was set > 40.

#### Supplementary Results

**Propionate induces intestinal mitochondrial protein hyper-propionylation in the context of high fat diet**

Protein propionylation in intestine was confirmed by western blots using an anti-propionyllysine antibody. The results showed that the level of propionylated protein in intestine collected from zebrafish fed HFSP0.5 diet was considerably higher than in that collected from zebrafish fed HFD (Supplementary Fig. 2A). The results of our study showed that Kpro in the HFSP0.5-zebrafish intestine occurred in the mitochondria (68%), cytoplasm (19%), extracellular space (10%), and cytoskeleton (3%) (Supplementary Fig. 2B). For a clear understanding of the biological role of Kpro, the GO of the Kpro proteins was annotated. Cell component analysis showed that the Kpro proteins were mostly localized in the mitochondria (Supplementary Fig. 2C), which is consistent with the subcellular distribution shown in Supplementary Fig. 2B. The main functions of propionyllysine are associated with the hydrogen ion transmembrane transporter activity and cation transmembrane transporter activity (Supplementary Fig. 2C). Biological processes of propionylated proteins are associated with the nucleoside metabolic process, nucleoside phosphate metabolic process and ribose phosphate metabolic process (Supplementary Fig. 2C). In the KEEG pathway analysis, genes related to the citrate cycle (TCA cycle) were highly enriched, including the protein malate dehydrogenase 2 (MDH2), citrate synthase (CS), isocitrate dehydrogenase (IDH), dihydrolipoyl dehydrogenase (DLDH) and succinate--CoA ligase (SUCLG2) (Supplementary Fig. 2C). PPI analysis was performed using STRING. A total of 19 mitochondria propionylated proteins were mapped to the PPI network after a cut-off at an interaction confidence of 0.7 (Supplementary Fig. 2D). As shown in Supplementary Fig. S2F, MDH2 and CS were highly interactive with other proteins.

**Efficiency of *si*RNA targeting *Sod2* and *Sirt3***

*Si*RNAs were used to knock down expression of *Sod2* and *Sirt3* in ZF4 cells. Scrambled *si*RNA was used as a negative control. *Sod2* was knocked down by 63.3% and 34.3% with *Sod2*-1 and *Sod2*-3, respectively (Supplementary Fig. 3A). Sirt3 was knocked down by 25.8%, 18.4% and 39.8% respectively (Supplementary Fig. 3B).

**The expression of *Pgc1α* and *Err* in response to propionate**

The expression of intestinal *Pgc1α* showed no significant difference between zebrafish fed HFD and HFSP0.5 diet (Supplementary Fig.4A). Exposure of OPP significantly elevated mRNA expression of *Sirt3* and *Err* in ZF4 cells (Supplementary Fig.4B-4D).

#### Supplementary Table

**Supplementary Table 1**. Ingredient and nutrient composition of experimental diets for one-month-old zebrafish (dry matter, g/kg).

| | one-month-old zebrafish | | | |
|---|---|---|---|---|
| Ingredients (g/kg dry diet) | LFD | LFSP0.5 | HFD | HFSP0.5 |
| Casein | 400 | 400 | 400 | 400 |
| Geltin | 100 | 100 | 100 | 100 |
| Wheat flour | 350 | 350 | 250 | 250 |
| Soybean oil | 60 | 60 | 80 | 80 |
| Lard oil | - | - | 80 | 80 |
| Sodium propionate[1] | - | 5 | - | 5 |
| Lysine | 3.3 | 3.3 | 3.3 | 3.3 |
| Ascorbyl phosphater | 1 | 1 | 1 | 1 |
| Vitamin premix[2] | 2 | 2 | 2 | 2 |
| Mineral premix[3] | 2 | 2 | 2 | 2 |
| Dicalcium phosphate | 20 | 20 | 20 | 20 |
| Choline chloride | 2 | 2 | 2 | 2 |
| Sodium alginate | 20 | 20 | 20 | 20 |
| Microcrystalline cellulose | 39.7 | 34.7 | 39.7 | 34.7 |
| Total | 1000 | 1000 | 1000 | 1000 |

[1]Sigma;

[2]Vitamin premix (g/kg): thiamine, 0.438; riboflavin, 0.632; pyridoxine·HCl, 0.908; *d*-pantothenic acid, 1.724; nicotinic acid, 4.583; biotin, 0.211; folic acid, 0.549; vitamin B-12, 0.001; inositol, 21.053; menadione sodium bisulfite, 0.889; retinyl acetate, 0.677; cholecalciferol, 0.116; *dl*-α-tocopherol-acetate, 12.632;

[3]Mineral premix (g/kg): $CoCl_2\cdot 6H_2O$, 0.074; $CuSO_4\cdot 5H_2O$, 2.5; $FeSO_4\cdot 7H_2O$, 73.2; NaCl, 40.0; $MgSO_4\cdot 7H_2O$, 284.0; $MnSO_4\cdot H_2O$, 6.50; KI, 0.68; $Na_2SeO_3$, 0.10; $ZnSO_4\cdot 7H_2O$, 131.93; cellulose, 501.09;

LFD, low-fat diet; LFSP0.5, low-fat diet supplemented with 0.5% sodium propionate; HFD, high-fat diet; HFSP0.5, high-fat diet supplemented with 0.5% sodium propionate.

**Supplementary Table 2**. Ingredient and nutrient composition of experimental diets for GF zebrafish larvae (dry matter, g/kg).

| | 5 dpf larvae | | |
|---|---|---|---|
| Ingredients (g/kg dry diet) | LFD | HFD | HFSP0.5 |
| Casein | 460 | 460 | 460 |
| Geltin | 110 | 110 | 110 |
| Wheat flour | 240 | 120 | 120 |
| Soybean oil | 35 | 80 | 80 |
| Lard oil | - | 80 | 80 |
| Sodium propionate[1] | - | - | 5 |
| Cod liver oil | 35 | 40 | 40 |
| Soybean ecithin | 20 | 20 | 20 |
| Lysine[8] | 3.7 | 3.7 | 3.7 |
| Ascorbyl phosphate | 1 | 1 | 1 |
| Vitamin premix[2] | 2 | 2 | 2 |
| Mineral premix[3] | 2 | 2 | 2 |
| Dicalcium phosphate | 20 | 20 | 20 |
| Choline chloride | 2 | 2 | 2 |
| Sodium alginate | 20 | 20 | 20 |
| Microcrystalline cellulose | 49.3 | 39.3 | 34.3 |
| Total | 1000 | 1000 | 1000 |

[1]Sigma, USA;

[2]Vitamin premix (g/kg): thiamine, 0.438; riboflavin, 0.632; pyridoxine·HCl, 0.908; *d*-pantothenic acid, 1.724; nicotinic acid, 4.583; biotin, 0.211; folic acid, 0.549; vitamin B-12, 0.001; inositol, 21.053; menadione sodium bisulfite, 0.889; retinyl acetate, 0.677; cholecalciferol, 0.116; *dl*-α-tocopherol-acetate, 12.632;

[3]Mineral premix (g/kg): $CoCl_2·6H_2O$, 0.074; $CuSO_4·5H_2O$, 2.5; $FeSO_4·7H_2O$, 73.2; NaCl, 40.0; $MgSO_4·7H_2O$, 284.0; $MnSO_4·H_2O$, 6.50; KI, 0.68; $Na_2SeO_3$, 0.10; $ZnSO_4·7H_2O$, 131.93; cellulose, 501.09;

LFD, low-fat diet; HFD, high-fat diet; HFSP0.5, high-fat diet supplemented with 0.5% sodium propionate.

**Supplementary Table 3**. *Si*RNA sequences.

| | Sense (5'-3') | Antisense (5'-3') |
|---|---|---|
| *Sod2*-1 | GCAGAGUCGGAUAUGUUCGTT | CGAACAUAUCCGACUCUGCTT |
| *Sod2*-2 | GCAAGCACCAUGCAACAUATT | UAUGUUGCAUGGUGCUUGCTT |
| *Sod2*-3 | GGCCAUAAAGCGUGACUUUTT | AAAGUCACGCUUUAUGGCCTT |
| *Sirt3*-1 | GCAGGCAUCAGCACACCAATT | UUGGUGUGCUGAUGCCUGCTT |
| *Sirt3*-2 | GCAGGAACAGUACCCAAAUTT | AUUUGGGUACUGUUCCUGCTT |
| *Sirt3*-3 | GCACUUCUUCACCUACCUUTT | AAGGUAGGUGAAGAAGUGCTT |
| Negative control | UUCUCCGAACGUGUCACGUTT | ACGUGACACGUUCGGAGAATT |

**Supplementary Table 4**. Quantitative PCR primers.

| Gene | Forward primer (5'–3') | Reverse primer (5'–3') |
|---|---|---|
| *Rps11* | ACAGAAATGCCCCTTCACTG | GCCTCTTCTCAAAACGGTTG |
| *Sirt3* | CATTAAATGTGGTGGAACAA<br>GAGGCCTG | AGTTCCTCTCCTTTGTAATCC<br>CTCCGAC |
| *Sod2* | ATGCTGTGCAGAGTCGGATA<br>TG | GCTGAAGGGAGACTTGGGTT |
| *Pgc1α* | CAGGTGGCAAGGCACAGAA<br>C | GCAGGACGACAGGGAGGAG<br>A |
| *Err* | TCAAAAGAACCATACAAGGC<br>AATA | CCACCAGCAGATGAGACACA<br>A |
| *Universal bacteria* | CCTACGGGAGGCAGCAG | ATTACCGCGGCTGCTGG |
| *Fusobacteria (phylum)* | KGGGCTCAACMCMGTATTGC<br>GT | TCGCGTTAGCTTGGGCGCTG |
| *Proteobacteria (phylum)* | TCGTCAGCTCGTGTYGTGA | CGTAAGGGCCATGATG |
| *Firmicutes (phylum)* | GGAGYATGTGGTTTAATTCG<br>AAGCA | AGCTGACGACAACCATGCAC |
| *Cetobacterium (genus)* | AGTTTGATCCTGGCTCAGGA<br>TG | GAGGCAAGTTCCTTACGCGT<br>T |
| *Plesiomonas (genus)* | CTCCGAATACCGTAGAGTGC<br>TATCC | CTCCCCTAGCCCAATAACAC<br>CTAAA |
| *Aeromonas (genus)* | GAAGGCCAAGTCGGCCGCC<br>AG | ATCTTGGCACGCCCGGGTTT<br>TC |

### Supplementary Figures

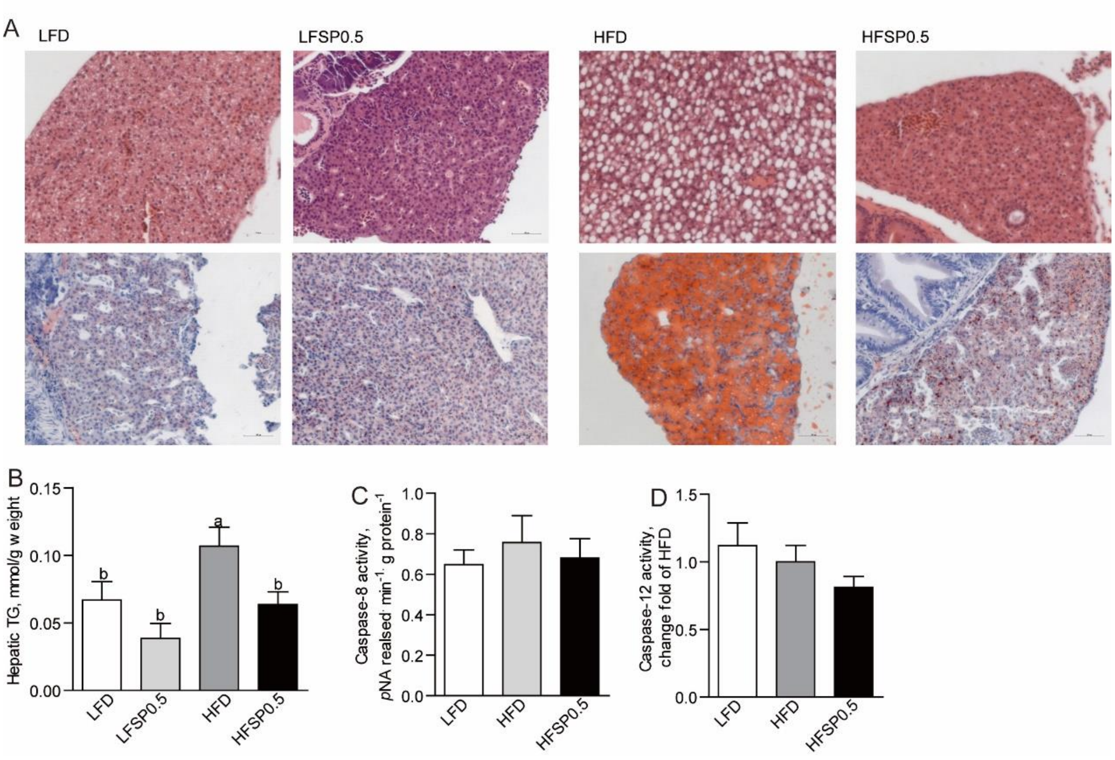


**Supplementary Fig. 1 Dietary propionate reduces hepatic fat accumulation in the context of HFD.** (A) Representative histopathologic image of H&E-stained and oil-red staining image in liver sections. (B) Hepatic TG in 1-month-old zebrafish fed LFD, LFSP0.5, HFD or HFSP0.5 diet for 2 wks. Intestinal (C) caspase-8 and (D) caspase-12 activity in 1-month-old zebrafish fed LFD, HFD or HFSP0.5 diet for 2 wks. Values are means ± SEMs (*n* = 6 biological replicates). Means without a common letter are significantly different, $P<0.05$. LFD, low-fat diet; HFD, high-fat diet; HFSP0.5, high-fat diet supplemented with 0.5% sodium propionate; LFSP0.5, low-fat diet supplemented with 0.5% sodium propionate.

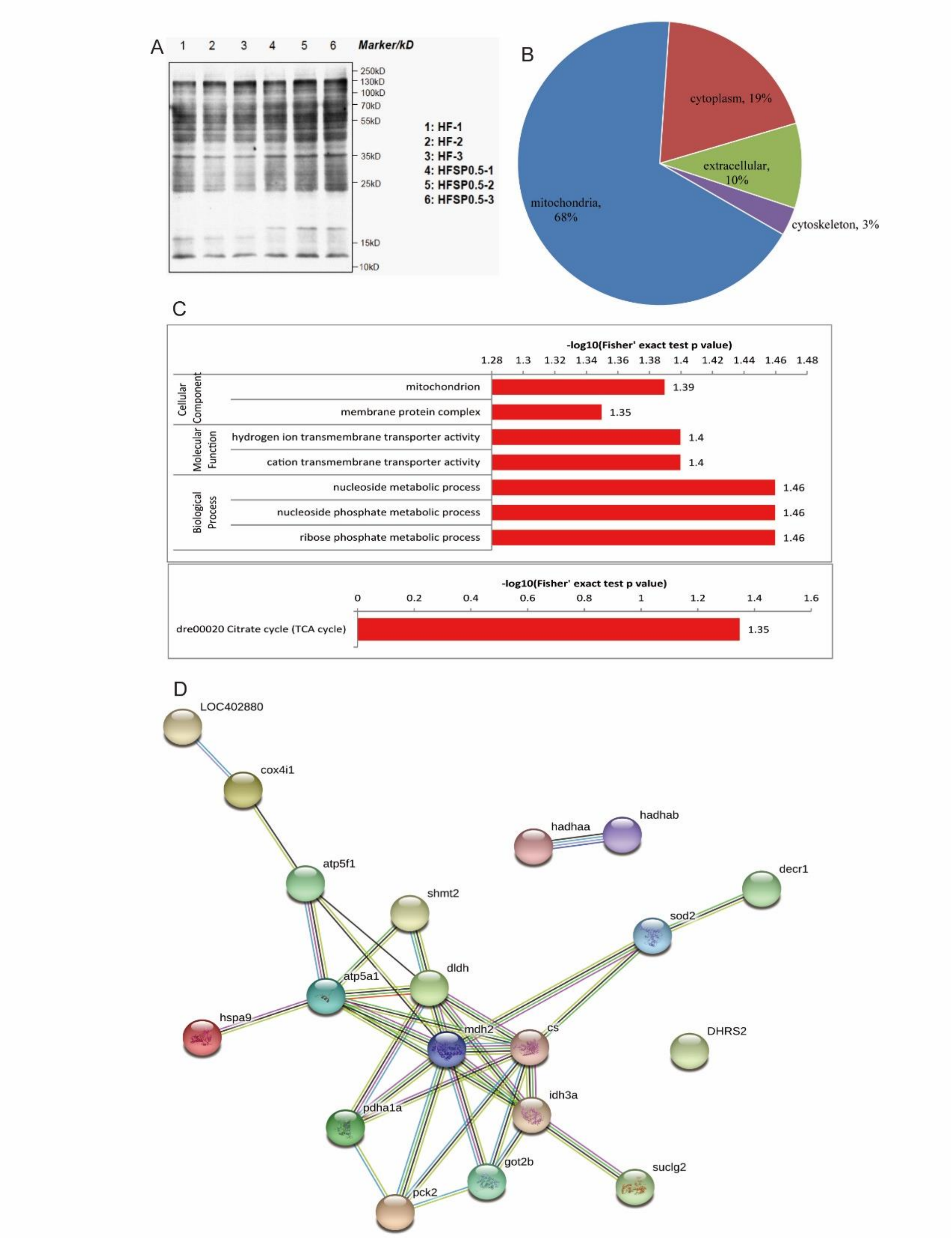


**Supplementary Fig. 2 Propionate induces intestinal mitochondrial protein hyper-propionylation in the context of high fat diet.** (A) Immunoblotting validation of Kpro from zebrafish intestine. Lanes 1-3: biological sample of HFD group, lanes 4-6: biological sample of HFSP0.5 group. (B) Subcellular distribution of increased lysine-propionylated proteins in zebrafish intestine of HFSP0.5 group compared to HFD group. (C) GO enrichment and KEEG pathway analysis of differentially represented Kpro proteins. (D) STRING analysis revealed mitochondrial Kpro protein interaction networks. Interactions of the identified Kpro proteins were mapped by searching the STRING database with a high confidence value of 0.7. HFD, high-fat diet; HFSP0.5, high-fat diet supplemented with 0.5% sodium propionate.

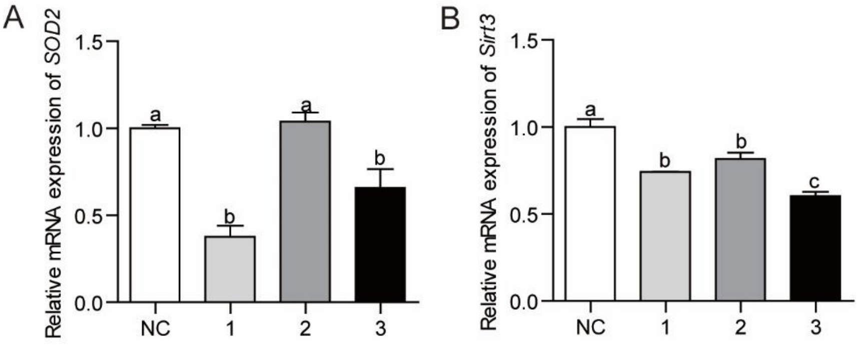


**Supplementary Fig. 3 Efficiency of *si*RNA knockdown.** (A) Efficiency of SOD2 *si*RNAs in ZF4 cells after transfection for 24 hrs. (B) Efficiency of SIRT3 *si*RNAs in ZF4 cells after transfection for 24 hrs. Values are means ± SEMs ($n$ = 3 biological replicates). Means without a common letter are significantly different, $P$<0.05. NC, negative control.

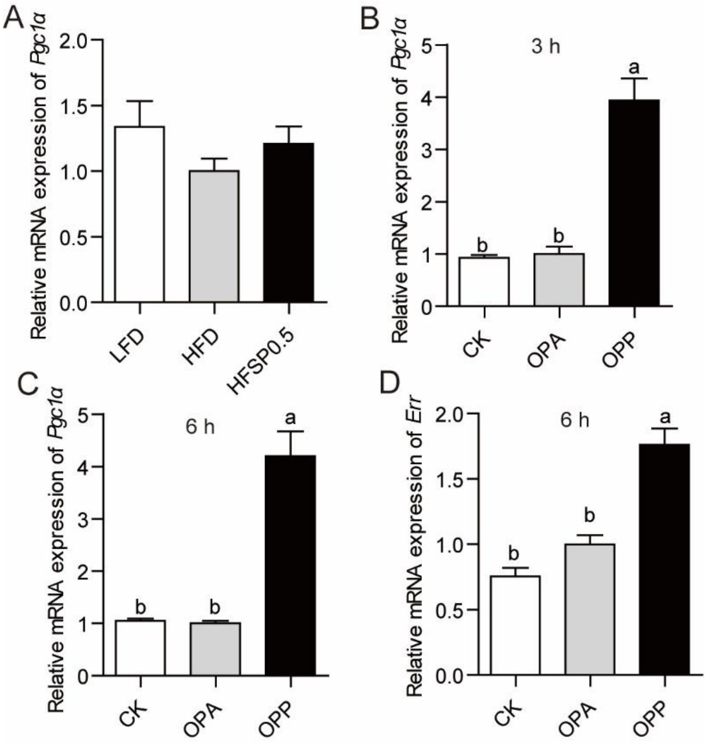


**Supplementary Fig. 4 The expression of Pgc1α and Err in response to propionate.** (A) The expression of intestinal *Pgc1α* in zebrafish fed HFD and HFSP0.5 diet. (B-C) The mRNA expression of *Pgc1α* in ZF4 cells treated with OPA or OPP. (D) The mRNA expression of *Err* in ZF4 cells treated with OPA or OPP. Values are means ± SEMs ($n$ = 4 or 6 biological replicates). Means without a common letter are significantly different, $P$<0.05. LFD, low-fat diet; HFD, high-fat diet; HFSP0.5, high-fat diet supplemented with 0.5% sodium propionate; OPA, mixture of 150 μM oleic acid and 50 μM palmitic acid; OPP, mixture of 150 μM oleic acid, 50 μM palmitic acid and 50 mM sodium propionate.
